# Self-Propelling Adaptive Robotic Microcatheters Enabled by Scalable Fabrication for Intracorporeal Navigation

**DOI:** 10.1101/2025.09.29.679200

**Authors:** Zhi Chen, Boris Rivkin, David Castellanos-Robles, Ivan Soldatov, Lukas Beyer, Mariana Medina Sánchez

**Affiliations:** NanoBiosystems Group, CIC nanoGUNE, San Sebastián, Spain; Institute for Solid State and Materials Research (IFW), Dresden, Germany; Center for Molecular Bioengineering (B CUBE), Dresden University of Technology, Dresden, Germany; Institute of Materials Science, Technische Universität Bergakademie Freiberg, Freiberg, Germany; IKERBASQUE, Basque Foundation for Science, Bilbao, Spain

**Author notes:** Corresponding author: Dr. Mariana Medina-Sánchez.

**Keywords:** magnetic soft continuum robot, microcatheter, microrobot platform, wave-crawling propulsion, minimally invasive surgery, intracorporeal navigation, continuous preparation

## Abstract

Minimally invasive therapies demand precise navigation through complex and delicate anatomical pathways, requiring medical tools that are small, flexible, and highly maneuverable. Here, we present a scalable fabrication platform for magnetic tubular microrobots, tethered and untethered, with programmable magnetization, enabling self-propulsion, and an adaptive remote control for targeted interventions. The platform uses Joule heating through a template wire for rapid, and reliable fabrication of microrobots with tunable dimensions. We demonstrate three device configurations: (1) a steerable guiding microcatheter with stiffness modulation; (2) an untethered tubular microrobot (TubeBot) exhibiting wave-crawling locomotion; and (3) a hybrid microcatheter robot that integrates distal-end wave-crawling propulsion with linear insertion to minimize tissue trauma. Validation in tortuous channels, soft phantoms replicating tissue compliance, 3D-printed organ models, ex vivo tissues, and live mice demonstrates the platform’s ability to achieve precise microrobotic navigation. The successful targeted delivery of sperm cells, embryos, and drug-mimicking compounds further highlights its potential for precision medicine, including applications in assisted reproduction and targeted drug delivery.

## 1. Introduction

In modern medicine, minimally invasive therapies are essential, providing reduced trauma, faster recovery, and fewer side effects.^[1]^ However, conventional tools for these procedures, such as laparoscopic instruments, guidewires, and catheters, typically range from millimeters to centimeters in diameter and often lack adaptive maneuverability, limiting their effectiveness, particularly in hard-to-reach regions of the body.^[2]^ Recent efforts have focused on optimizing these devices for mechanical performance, miniaturization, and functionality, while minimizing tissue damage.^[3]^ For instance, in reproductive medicine, falloposcopy has made significant advances in miniaturization, enabling direct visualization of the fallopian tubes using tiny flexible endoscopes. This allows detection of tubal abnormalities, assessment of the mucosal lining, and the performance of minor therapeutic interventions, such as clearing small obstructions.^[4]^ However, these devices remain largely passive, relying on mechanical pushing from outside the body. They tend to scrape the fallopian tube cilia during insertion, and they face challenges navigating the complex and variable anatomy of the fallopian tubes (See Table S1 in the Supporting Information for a comparative overview). Although fluoroscopy holds therapeutic potential, its use is primarily limited to diagnostic purposes, typically in cases where hysterosalpingography (HSG) is insufficient. HSG, a radiologic technique in which contrast dye is injected into the uterus and fallopian tubes via a catheter, is less invasive in principle but requires radiation exposure for imaging which may alter reproductive physiology.^[5]^ Similarly, procedures such as artificial insemination near the fertilization site (sperm sample release), intrafallopian embryo transfer, and endometriosis/cancer management often rely on laparoscopy, which requires multiple instruments for imaging, illumination, and surgical intervention.^[6]^ These tools are relatively large and can increase the risk of tissue injury, infection, or, in some cases, ectopic pregnancies.^[6a, 7]^ Comparable challenges exist in other biomedical scenarios such as the peripheral microcirculation or terminal bronchioles and alveolar sacs in the lungs.^[2c, 8]^

In this context, continuum robots (CRs) offer a promising solution. Their structures bend continuously along their length, enabling navigation through tortuous anatomical pathways while remaining connected to a control hardware.^[9]^ CRs have been applied successfully in clinical procedures including cardiac ablation, lung biopsy, and transoral robotic surgery.^[10]^ But they are still large in size (mm to cm range), and typically rely on tendon-like, hydraulic, shape-memory alloy, or polymer-based mechanisms for distal-tip manipulation (Table S2).^[11]^ An strategy to miniaturize the CRs further, is by incorporating magnetic materials into flexible segments to enable their remote actuation using global magnetic fields.^[12]^ Replacing rigid permanent magnets with micro- or nanoparticles further alleviates size constraints but their weaker magnetic moments require stronger external fields to achieve effective actuation.^[13]^ These challenges highlight the need for novel actuation strategies to enhance navigational performance in intricated organs.^[14]^ Untethered microrobots provide a possible solution, enabling remote actuation at ultra-small scales (Table S3).^[15]^ Potential applications include assisting sperm with motility deficiencies in reaching the oocyte for in vivo assisted reproduction, targeted cancer therapy, management of infectious diseases, and tissue engineering, among others.^[16]^ Nonetheless, concerns remain regarding residual materials left in the body, as their retrieval or degradation after therapy is often challenging.^[17]^ Some studies have demonstrated combining catheters with untethered microrobots to enable their controlled release and retrieval. For instance, a flexible catheter equipped with a miniaturized electromagnet allowed precise release and localized actuation of microrobots deep within the body.^[18]^ Similarly, clinical endoscopes have been employed to deliver magnetic cell-based robots for high-precision stem cell therapy.^[19]^ Magnetically guided microcatheters have been reported to inject swarms of magnetic particles into the circulatory system.^[20]^ Another strategy utilizes multi-segment catheters linked by biodegradable connectors, allowing the controlled release of untethered agents for precise interventions.^[21]^ And catheters incorporating permanent magnets have been used to retrieve nanoscale agents from the bloodstream.^[22]^ Despite the translational potential of these technologies, significant limitations persist, including the size constraints of electromagnet-equipped catheters and endoscopes, restricted autonomous navigation, and difficulties in achieving scalable and reproducible fabrication.

Therefore, we suggest a fabrication approach for mass-producing tubular magnetic microrobots, both tethered and untethered, that can be magnetized and operated across diverse microenvironments (Figure 1A). These tubular structures feature lumen microchannels capable of housing drug-loaded, stimuli-responsive hydrogels or enabling the capture and release of cells and other therapeutic cargo. Inspired by the wave-crawling propulsion observed in certain untethered microrobots, we introduce a hybrid motion strategy to enhance maneuverability: the distal portion of the microcatheter self-propels while being gently pushed during insertion. The structures are fabricated using Joule heating with a wire template, allowing magnetization either during or after production (Figure 1 and Video S1). We present three configurations: (1) a steerable guiding microcatheter with tunable stiffness; (2) a wave-crawling tubular untethered microrobot (TubeBot) incorporating an enzyme-digestible hydrogel loaded with a model drug; and (3) a hybrid microcatheter robot combining distal wave-crawling propulsion with linear insertion. The platform is validated by demonstrating navigation and cargo delivery across different models, including a soft tissue phantom, a 3D tortuous channel, a full-scale human bronchial tree, 2D and 3D reproductive organ models, ex vivo mouse reproductive and respiratory organs, and an in vivo mouse model. Furthermore, we demonstrate targeted sperm delivery near the ampulla of the fallopian tube, for enabling artificial insemination, which is appealing in cases of male infertility such as oligospermia (low sperm count) or asthenospermia (low sperm motility).^[23]^ In this work we also show the intrafallopian embryo transfer, relevant for cases of recurrent embryo implantation failure, which under in vivo conditions, can develop under natural, nutrient-rich environment synchronized with endometrial preparation.^[17b, 24]^ Beyond reproductive medicine, we demonstrate that magnetic microcatheters can access distal lung regions for targeted therapy, aiming to enhance drug delivery to the alveoli or bronchioles, in contrast to conventional lung-targeted therapies, which are typically administered systemically via intravenous injection, which is less precise and subject to hepatic metabolism. The methods and devices presented here, along with the supporting demonstrations, establish a solid foundation for biologically and clinically relevant studies. This paves the way for the platform’s translational application in precision medicine, particularly in reproductive medicine, with potential for expansion into other biomedical fields.

**Figure 1.**
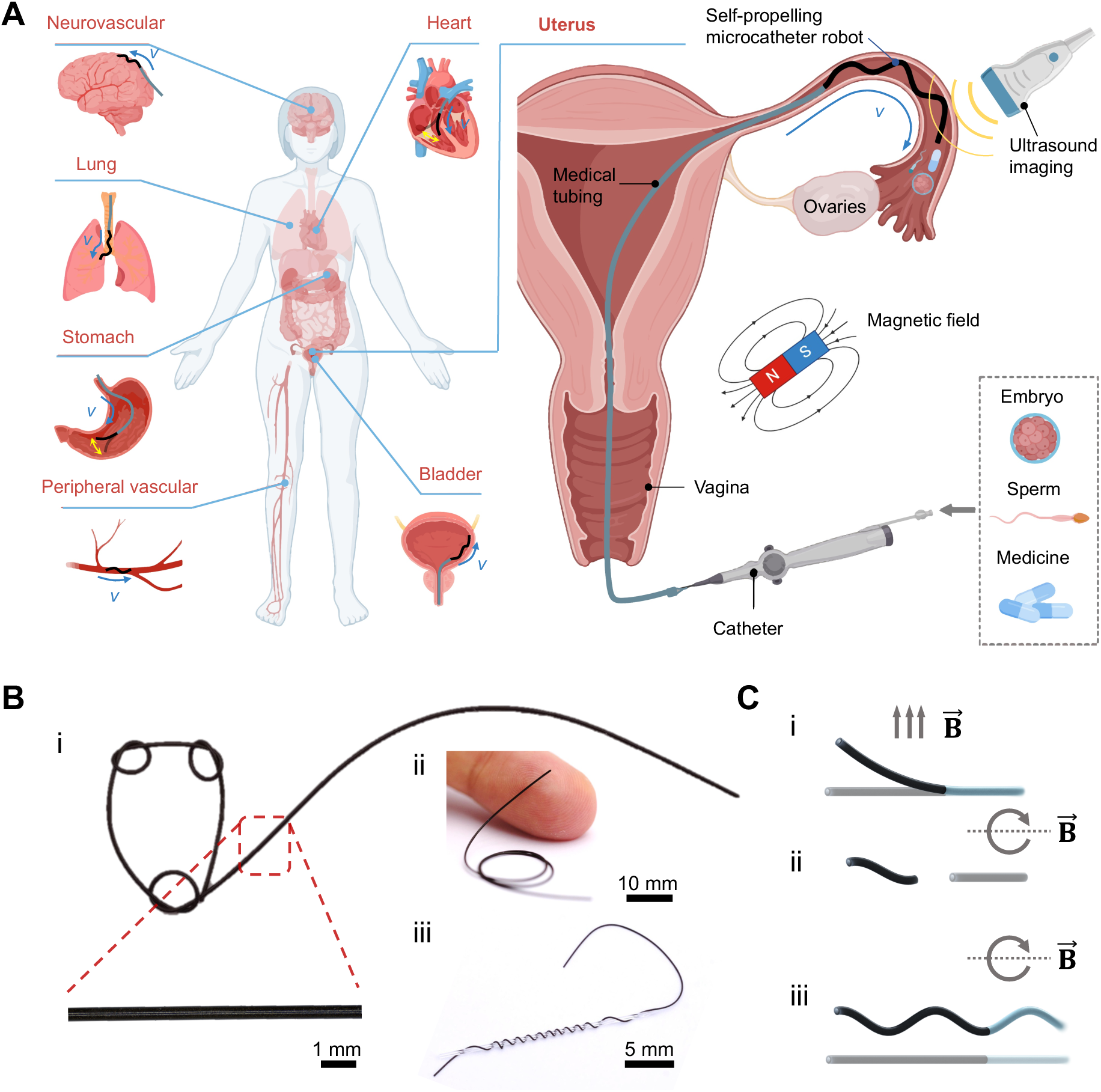
Schematic illustration of a robotic microcatheter platform. (A) The microcatheter platform is designed for accessing difficult-to-reach areas within the human body and can be configured in various ways depending on the specific application. (B) The microcatheter, with an outer diameter of 400 μm, is soft and flexible, demonstrating high compliance as shown when wrapped around a 1 mm diameter capillary. (C) The platform includes various device configurations, such as (i) a guiding microcatheter, (ii) a tubular microrobot, and (iii) the microcatheter robot.

## 2. Results and Discussion

### 2.1. Microcatheter Preparation and Performance

To enable rapid, low-cost fabrication of magnetic microcatheters with micron-scale diameters, we employed Joule heating to solidify a resin mixture through a continuous curing process (**Figure 2A**(i) and Video S1). Ferromagnetic materials such as NdFeB (neodymium-iron-boron) exhibit strong induced magnetization under an applied magnetic field. Unlike soft magnetic materials, which easily lose magnetization once the external field is removed, hard magnetic materials like NdFeB possess high coercivity, allowing them to retain significant remanent magnetization after the magnetic field is removed.^[20]^ The resin mixture consisted of polydimethylsiloxane (PDMS) resin (mixed at a 10:1 ratio) and uniformly dispersed NdFeB particles with a diameter of 5 μm.

**Figure 2.**
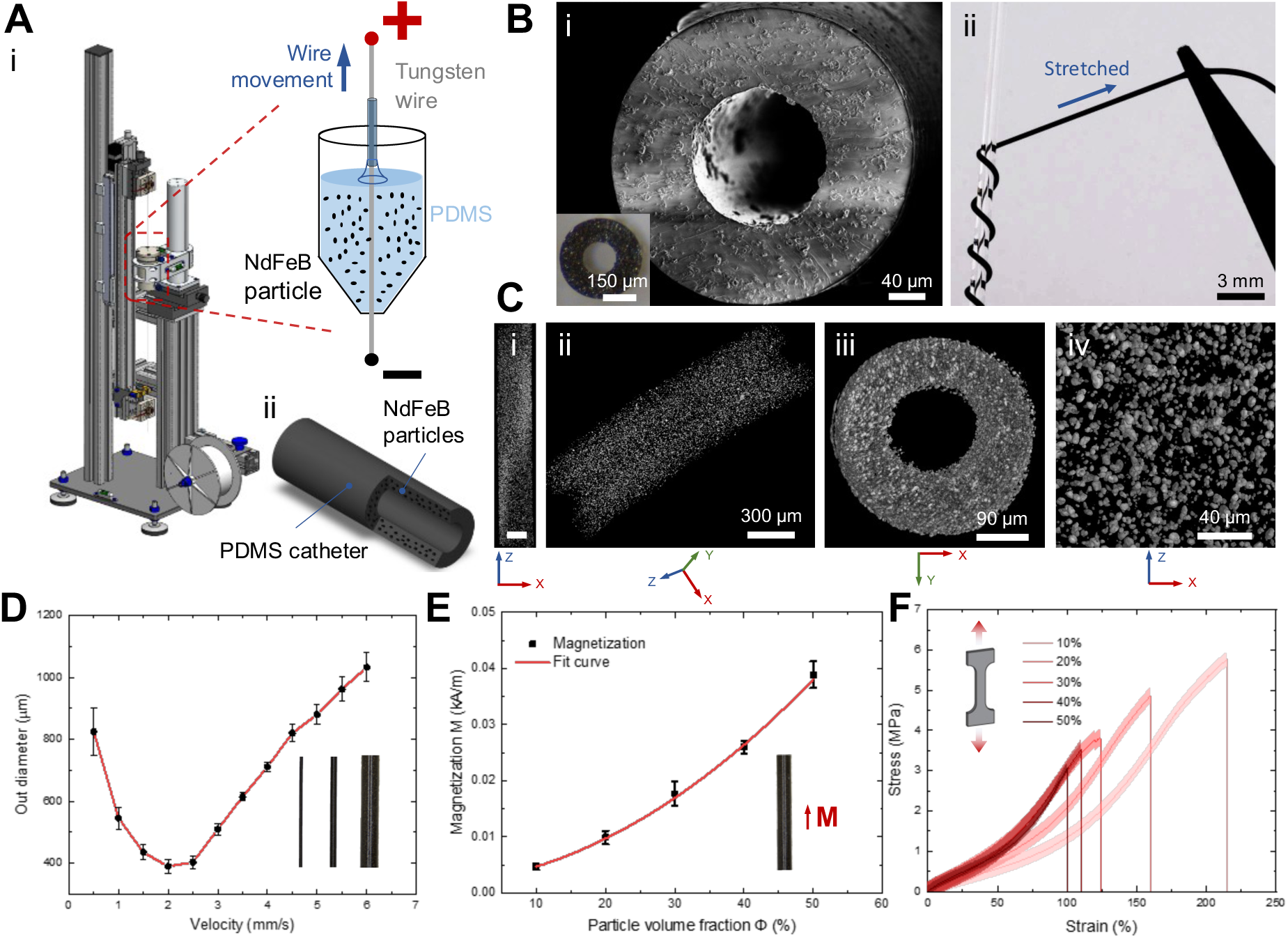
Fabrication and characterization of the microcatheter platform. (A) The fabrication setup, detailing the composite composition and the expected resulting geometry. (B) (i) SEM image of the microcatheter cross-section and (ii) a photograph of the microcatheter under tension. (C) MicroCT images show the distribution of NdFeB magnetic particles in the microcatheter at different magnifications, including longitudinal view (i) (scale bar: 200 μm), oblique view (ii), cross-sectional view (iii), and local magnified view (iv). (D) Control over the microcatheter diameter using the Joule heating principle, with variation achieved by adjusting the tungsten wire pulling speed. (E) Magnetization (*M*) of the microcatheter as a function of the concentration of embedded magnetic particles (φ). (F) Stress-strain curves of ferromagnetic composites at different magnetic particle concentrations.

The microcatheter fabrication setup is depicted in Figure S1. The resin mixture was placed in a central container with tungsten wire running through its center, fixed at both ends to a displacement stage. A direct current was applied to the tungsten wire to generate heat, initiating the solidification of the resin mixture adhered to the wire. As the displacement stage gradually pulled the heated tungsten wire upward, the viscous resin coated the wire uniformly and solidified, forming the microcatheter. After manually removing the tungsten wire, a magnetic microcatheter was obtained, with its length determined by the tungsten wire’s length. The structure of the fabricated microcatheter is shown in Figure 2A(ii), where NdFeB particles are uniformly embedded within the cured PDMS.

Scanning electron microscopy (SEM) characterization of the microcatheter cross-section (Figure 2B(i)) revealed a centrally located working channel, with a uniform and crack-free wall structure. To further examine the internal distribution of NdFeB particles, X-ray computed tomography was employed to obtain a three-dimensional representation of the particle distribution (Figure 2C). Since the PDMS matrix signal is challenging to distinguish from NdFeB, segmentation based on particle size observed in SEM images (Figure S2) was performed to approximate the true three-dimensional distribution of particles. The results indicated that the NdFeB particles were homogeneously distributed throughout the PDMS matrix with minimal overlap, confirming the uniformity of the microstructure.

The inner diameter of the microcatheter was determined by the diameter of the tungsten wire, which remained unchanged during manual removal. With a constant power input to the tungsten wire, the outer diameter of the microcatheter was influenced by the speed at which the wire was pulled during heating (Figure 2D). As the pulling speed increased from 0 mm/s to approximately 2 mm/s, the outer diameter gradually decreased, reaching a minimum of around 350-400 μm. When the pulling speed increased further to 6 mm/s, the outer diameter began to rise again, reaching approximately 1000 μm at the maximum speed tested. Thus, the outer diameter of the fabricated microcatheters ranged from 350 to 1000 μm. In the range of 0.5-2 mm/s, the tungsten wire moves very slowly at a speed of 0.5 mm/s. The tungsten wire stays in the resin mixture for a longer time, so the curing effect is noticeable, and a microcatheter with an outer diameter of about 800 μm can be prepared. As the pulling speed increases, the residence time of the tungsten wire in the resin mixture decreases, and because the gravity effect of the resin mixture is significant, the resin attached to the surface of the tungsten wire decreases, and the outer diameter of the prepared microcatheter decreases. When the tungsten wire pulling speed is in the range of 2-6 mm/s, the outer diameter of the microcatheter is mainly affected by the viscosity of the resin mixture, which increases with the increase in the pulling speed. The increasing diameter at increasing speeds is a consequence of a hydro-dynamic phenomenon, described by the theory of Landau-Levich-Derjaguin, as elaborated on in the “Landau-Levich-Derjaguin theory” section of the Supporting Information. Therefore, the outer diameter of the microcatheter prepared by this rapid and continuous method can be between 0.35 and 1 mm, and the diameter of the microcatheter can be flexibly selected according to different usage scenarios.

The magnetic and mechanical properties of the microcatheter varied with the NdFeB particle content. Microcatheters with five different mass fractions of NdFeB particles were fabricated and characterized for magnetization (Figure 2E). The magnetization increased with particle concentration but did not follow a strictly linear relationship. Vibrating sample magnetometry (VSM) tests conducted on ferromagnetic composite resin cured at different temperatures revealed noticeable differences in remanent magnetization in the **M**(H) curves (Figure S3). The microstructure of NdFeB particles near the tungsten wire is changed by the transient high temperature, which enhances the soft magnetic properties of NdFeB particles.^[21]^ However, since the overall curing time was short, most NdFeB particles still retain their hard magnetic properties.

Dog-bone-shaped specimens were fabricated for tensile testing using templates.^[12]^ Mechanical properties of the specimens with varying particle concentrations were obtained through tensile testing (Figure 2F). The results showed that the stress-strain behavior was consistent across different specimens, with the sample containing a 10% mass fraction of NdFeB exhibiting the highest tensile strain, while the 50% particle content sample showed the lowest tensile strain. Therefore, increasing the NdFeB particle content led to increased stiffness of the microcatheter, although even at 50% loading, the microcatheter retained 100% tensile deformation capacity.

### 2.2. The Guiding Microcatheter

The embedded NdFeB particles within the microcatheter were magnetized using a strong uniform axial magnetic field (***B*_1_ = 3 T**), resulting in a consistent magnetic moment (***m***) along the length of the microcatheter, thereby producing a guiding microcatheter (**Figure 3A**). When an external uniform magnetic field (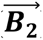) was applied perpendicularly to the axis, the tip of the guiding microcatheter attempted to align with the field direction, resulting in the overall bending of the catheter. When coupled with medical tubing, the guiding microcatheter could achieve effective steering and navigation under the influence of a uniform magnetic field.

**Figure 3.**
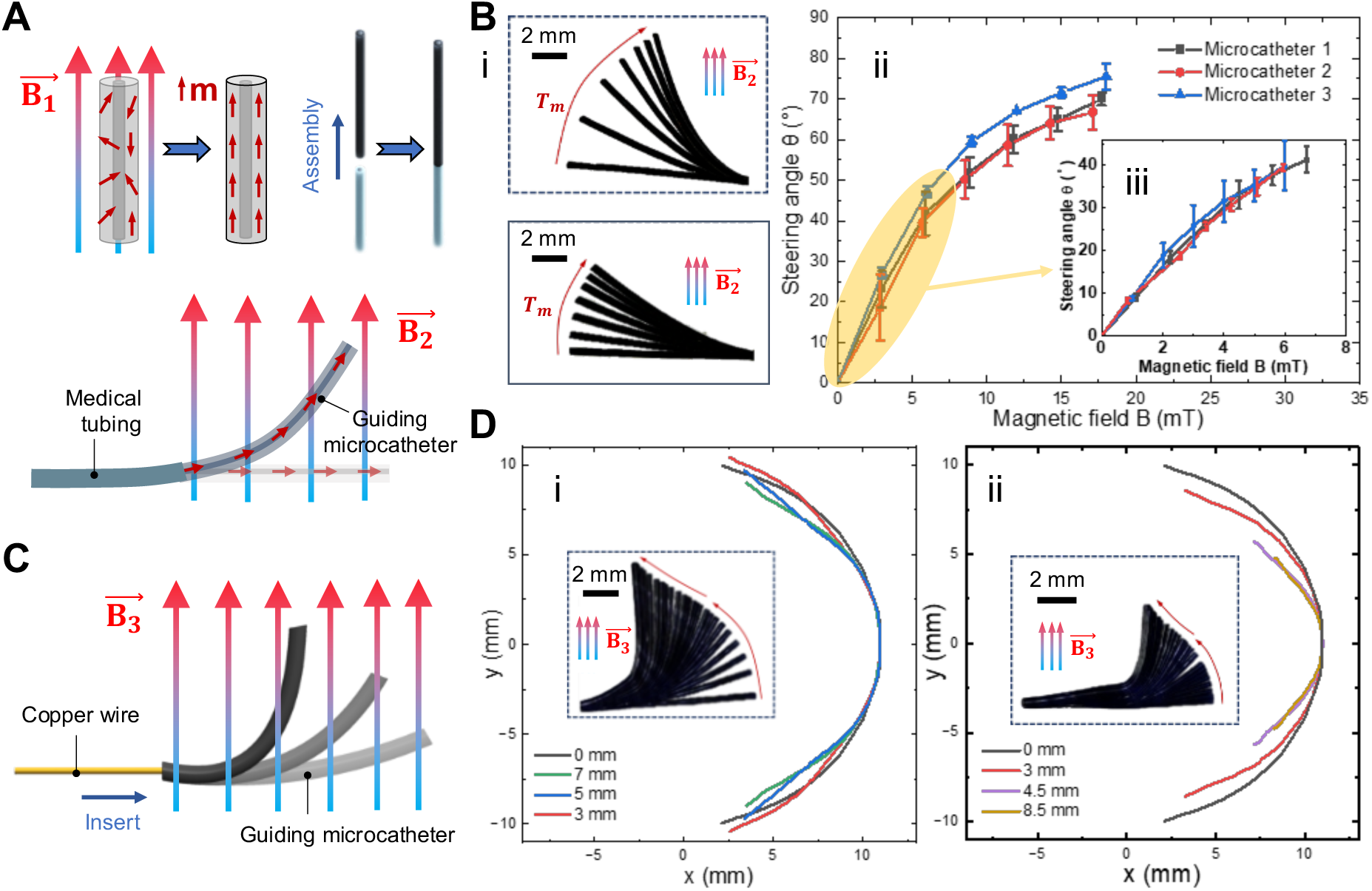
Characterization of the guiding microcatheter. (A) Schematic diagram of a microcatheter magnetized internal magnetic moment m with a uniform magnetic field 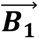 along the axial direction. It’s toward the direction of the uniform magnetic field 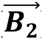 applied perpendicularly to the body. (B) (i) The microcatheter achieves different angles (*θ*) of deflection due to the magnetic torque *T_m_* in the uniform magnetic field 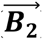. The magnetic field strength range is 0 to 18 mT (ii), and the steering angle increases with the magnetic field strength, especially in the magnetic field strength range is 0 to 7 mT, *θ* increases nearly linearly (iii). (C) Schematic diagram of inserting the copper wire into the guiding microcatheter to change the local stiffness of the guiding microcatheter. (D) Distribution of catheter-Cu-wire assembly with different diameters of Cu wire due to the guidance of a uniform magnetic field 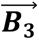 (0 to 100 mT). (i) Catheter-Cu-wire assembly of different lengths and 50 μm diameter, (ii) 100 μm diameter.

To assess its steering capability, the guiding microcatheter was suspended in the magnetic field, demonstrating sufficient magnetic torque to overcome gravity in fields ranging from 0 to 18 mT (Figure 3B(i)). Further characterization involved testing the steering angles of guiding microcatheters with the same diameter and length in different magnetic field strengths (Figure 3B(ii) and (iii)). The results showed that the steering angle could reach up to 80° at a field strength of 18 mT. Although the change in steering angle was not directly proportional to the applied magnetic field strength, the results demonstrated reproducibility and the ability of the guiding microcatheter to achieve significant steering at relatively low field strengths. At magnetic field strengths within 7 mT, the change in the steering angle is approximately linearly related to the applied magnetic field. This provides a way to precisely control the small-angle (0 to 40°) deflection of the guiding microcatheter at low magnetic field strengths.

The magnetic steering and selective navigation capabilities of the guiding microcatheter were further validated using a two-dimensional maze (Figure S4 and Video S2). The maze was designed with multiple branches (angles ranging from 0° to 50°) and a channel width of 1.5 mm. Magnetic steering was achieved using an external permanent magnet, which applied the magnetic torque needed for turning, while manual pushing was used to propel the medical tubing forward.

Since the steering and workspace are dependent on the tip position of the guiding microcatheter, a fixed length limits the workspace to a shell volume. Altering the workspace requires changing the length of the guiding microcatheter in free space, which imposes stringent requirements on both operations and in vivo space.^[22]^ To address this, materials were inserted or infused into the microcatheter’s central channel to locally alter stiffness, thereby enabling the tip to reach various positions within the body cavity without changing the overall length of the guiding microcatheter. Copper wires were chosen for insertion into the working channel to form stiffness difference (Figure 3C). By adjusting the length of the inserted copper wire, various tip positions were achieved under the influence of an external magnetic field (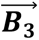, 0-100 mT).

Copper wires with diameters of 50 μm (Figure 3D(i)) and 100 μm (Figure 3D(ii)) were inserted into the catheter, and the deflection of the guiding microcatheter tip was measured for different insertion lengths. The tip achieved a deflection of up to 90°, and its trajectory was recorded throughout the deflection process. When a magnetic field was applied with copper wire inserted, the tip trajectory differed from the one without wire (0 mm insertion). Increasing the length of the copper wire reduced the tip’s trajectory radius, which was more pronounced when using the 100 μm wire (e.g., the 3 mm insertion case for both 50 μm and 100 μm wires). This indicates that steering and workspace adjustments can be achieved in more confined spaces.

The observed effect is attributed to the difference in stiffness between the copper wire and the guiding microcatheter. Increasing the length of the copper wire resulted in greater stiffness of the catheter segment containing the wire, leading to a smaller bending angle under the magnetic field. In contrast, the segment without the copper wire retained its flexibility, allowing for larger deflections. The tip position was thus influenced by the combined effects of both stiffened and flexible segments. Using a stiffer copper wire further enhanced the guiding microcatheter’s ability to navigate confined spaces (Figure 3D(ii)).

In addition, to more clearly demonstrate the distribution of the working space of the guiding microcatheter in a uniform magnetic field due to the difference in stiffness (Figure S5), copper wires of different lengths with a diameter of 150 μm were inserted into the working channel. By using the built-in wire with higher stiffness, the flexible length of the guiding microcatheter was changed, thereby achieving large-angle navigation in a narrower space. However, we also noticed that this application still has limitations, because deep in the anatomical structure, it is difficult to completely avoid wire puncture of the guiding microcatheter to deliver high-stiffness wires to the guiding microcatheter to form a stiffness difference. Although no similar problems occurred during our testing, it is still noteworthy.

### 2.3. The Tubular Microrobot (TubeBot)

To navigate complex pathways, tubular microrobots need to generate undulatory body waves within confined spaces to enable crawling propulsion.^[23]^ To achieve this in a magnetic field, the microcatheter required a sinusoidal magnetization along its length. A microcatheter segment was cut to 4 mm in length and fixed in a cylindrical mold with a radius of 0.5 mm for magnetization (**Figure 4A**(i)). A uniform magnetic field (***B*_1_ = 3 T**) was applied perpendicularly through the mold and microcatheter, ensuring an initial phase shift of 45° for the magnetization (Figure 4A(i), ***α*** = 45°), resulting in a tubular microrobot.^[24]^

**Figure 4.**
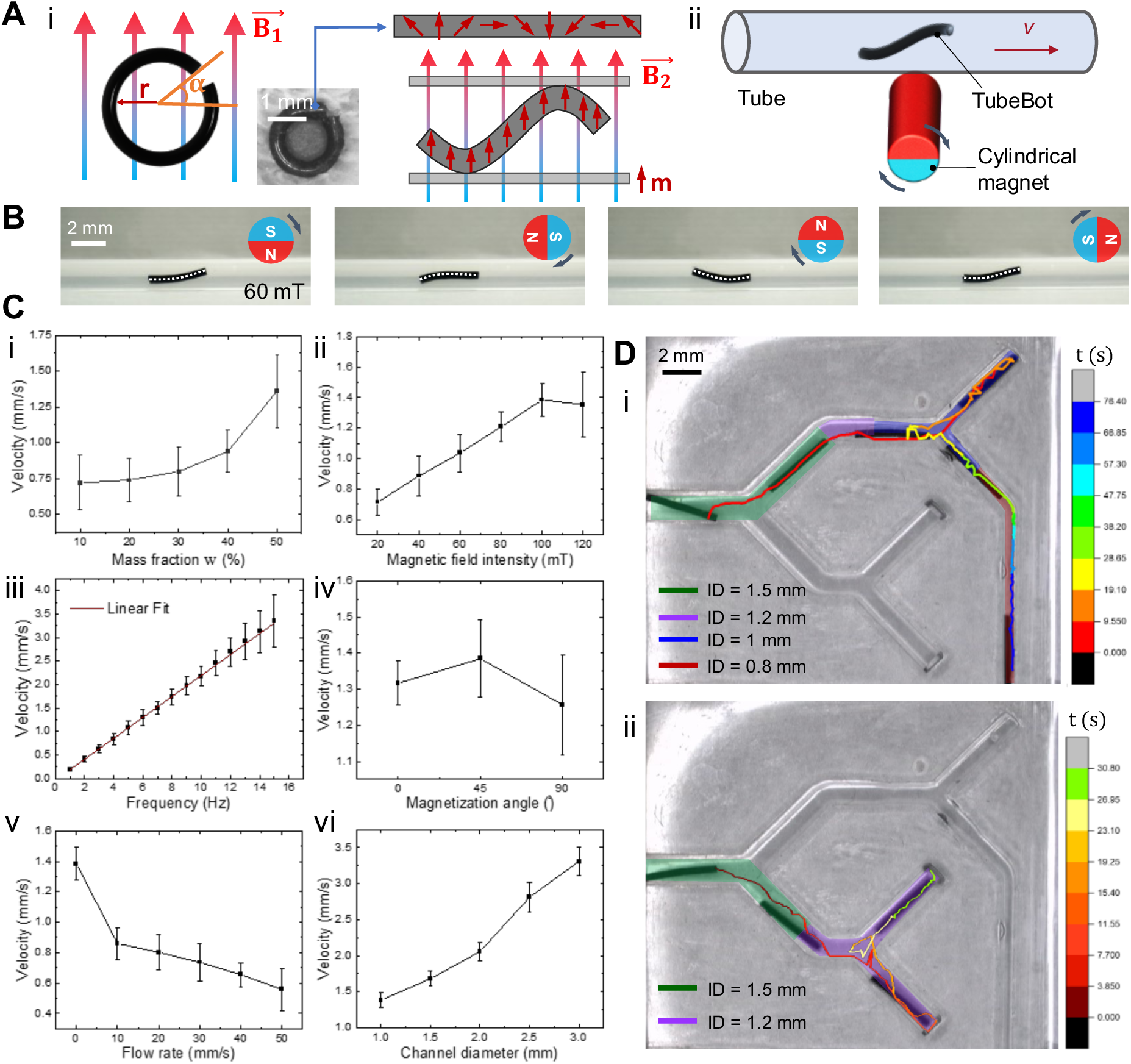
Tubular microrobot (TubeBot) characterization. (A) (i) The TubeBot is fixed in a mold (radius = 1 mm) and initially magnetized using a uniform magnetic field 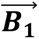. After magnetization, the free state of TubeBot is a straight line. When a uniform upward magnetic field 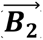 is applied, TubeBot exhibits a wavy shape in a narrow space. (ii) Schematic diagram of the test device for TubeBot. (B) Postures of TubeBot at different angles within a rotating magnetic field inside a phantom silicone tube (inner diameter = 1 mm). (C) Factors affecting crawling speed of the TubeBot: (i) increased magnetic particle content, (ii) stronger magnetic field amplitude, (iii) higher rotation frequency of the magnetic field, (iv) variation with magnetization angle, peaking at 45°, (v) when the TubeBot is against fluid, the crawling speed of the microcatheter robot is linearly related to the fluid flow rate. (vi) The crawling speed of the TubeBot in silicone tube of different diameters increases as the tube diameter increases. (D) TubeBot navigating and crawling in a channel with variable diameter, with an image overlay showing the microrobot at different positions while moving along the channels. (i) TubeBot navigating in path 1, with one end tracked to record movement over time. (ii) TubeBot navigating in path 2.

To characterize the programmed magnetization, a magneto-optical Kerr effect (MOKE) microscope was used to image the magnetic distribution within TubeBot (Figure S6). The alternating bright and dark areas with 2 nodes along the microcatheter, confirmed that the internal magnetic distribution was consistent with the design, demonstrating the feasibility of magnetization programming. Without an external field, TubeBot remained a straight horizontal line. When a uniform magnetic field (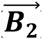) was applied perpendicular to the TubeBot’s body axis, the embedded NdFeB particles attempted to align with the external field, generating magnetic torque and causing deformation. When placed in a confined space and subjected to a rotating magnetic field, the shape of TubeBot becomes sinusoidal, enabling crawling propulsion.

To evaluate TubeBot’s propulsion within a vessel-like environment, a circular silicone tube filled with phosphate-buffered saline (PBS) solution was used, and a cylindrical permanent magnet generated the rotating magnetic field (Figure 4A(ii)). A rotating magnetic field was applied to TubeBot in a tube with an inner diameter of 1 mm, and its deformation over one rotation cycle (***B*_2_ = 60 mT**) was recorded (Figure 4B). Due to the body length not being significantly greater than the diameter, TubeBot did not exhibit a complete sinusoidal shape (the white dashed line represents TubeBot’s posture).^[24]^ Testing the crawling direction of TubeBot revealed that it moved in the same direction as the magnetic field rotation (Figure S7). To optimize TubeBot’s crawling performance, various types of TubeBots, magnetic field parameters, and propulsion environments were studied (Figure 4C). The effect of embedded magnetic particle content on TubeBot’s crawling speed was evaluated by measuring the crawling speeds of TubeBots with different mass fractions of magnetic particles (***w***) under a magnetic field of 5 Hz and 60 mT. As shown in Figure 4C(i), the crawling speed increased significantly as **w** increased. For mass fractions between 10% and 30%, the speed remained around 0.75 mm/s with little variation. However, at **w** greater than 30%, the speed increased markedly, reaching approximately 1.5 mm/s at ***w*** = 50%. Higher magnetic particle content significantly increased TubeBot’s stiffness, resulting in reduced deformation under the magnetic field and unstable crawling speed, thus limiting further increases in particle content.

TubeBot’s crawling speed under different rotating magnetic field strengths (Figure 4C(ii)) showed an approximately linear increase, ranging from about 0.7 mm/s to 1.5 mm/s, as the amplitude of the field increased from 20 mT to 100 mT, with the magnetic field rotation frequency held at 5 Hz. However, a slight decrease in crawling speed was observed when the field strength reached 120 mT. This reduction could be attributed to increased deformation of TubeBot at higher magnetic field strengths, which led to changes in its crawling posture, thereby affecting the crawling performance. Additionally, at higher magnetic field strengths, the attraction of the rotating permanent magnet to TubeBot increased, further impeding its crawling speed and introducing greater variability.

The crawling speed of TubeBot under different magnetic field rotation frequencies (Figure 4C(iii)) exhibited a linear growth trend as the frequency increased from 2 Hz to 16 Hz, with a magnetic field strength of 100 mT. The crawling speed increased from approximately 0.25 mm/s to 3.25 mm/s, and the experimental data showed good agreement with the linear fit, indicating a direct correlation between rotation frequency and crawling speed.

Based on the exploration of the effect of external magnetic parameters on TubeBot’s crawling performance and for the convenience of data collection, subsequent characterization of TubeBot’s motion—including factors such as magnetization angle, resistance to fluid, varying tube diameters, magnetization angle, magnetization diameter, and robot length—was conducted using a rotating magnetic field of 5 Hz and 100 mT, unless otherwise stated.

The crawling speed of TubeBot is strongly influenced by its magnetization angle, **α**, which is defined as the angle between one end of the TubeBot and the direction of the applied magnetic field (Figure 4A(i)). The magnetization angle affects the initial phase shift of the internal magnetization distribution, thereby altering the TubeBot’s crawling posture and speed. As shown in Figure 4C(iv), the crawling speed reached its peak when the initial magnetization angle was set to 45°. This suggests that a 45° magnetization angle provides an internal magnetization distribution that most closely resembles a sinusoidal wave, resulting in more efficient crawling.

The ability of TubeBot to move within fluid-filled environments was assessed by testing its performance in a silicone tube with various flow rates of PBS (Figure 4C(v)). TubeBot crawled against the direction of the PBS flow, and the flow velocity was controlled using a micropump. Initially, TubeBot’s crawling speed decreased rapidly, followed by a more gradual decline as the PBS flow rate increased. At a flow rate of 50 mm/s, TubeBot maintained a net forward speed of 0.6 mm/s. For reference, venous blood flow in human leg veins has a peak velocity of 200-300 mm/s and a mean velocity of 20-40 mm/s.^[25]^ Thus, TubeBot demonstrated the potential to crawl against venous blood flow in vessels.

In addition to straight pathways, in vivo channels are typically non-uniform in diameter. Therefore, TubeBot’s performance in different channel diameters was evaluated by measuring its crawling speed in round channels of varying diameters (Figure 4C(vi)). As the channel diameter increased from 1.0 mm to 3.0 mm, TubeBot’s crawling speed showed an approximately linear increase, from about 1.35 mm/s to 3.25 mm/s. This result indicates that TubeBot achieved higher crawling efficiency in larger-diameter channels. Moreover, we investigated the influence of different magnetization diameters and TubeBot lengths on crawling speed (Figure S8). The results indicated that variations in magnetization diameter led to asymmetric magnetization within TubeBot of the same length, resulting in different left-right crawling speeds. The relationship between TubeBot length and crawling speed was approximately linear, with longer TubeBots exhibiting decreased speed. TubeBots for various testing requirements were magnetized following different schemes, as illustrated in Figure S9.

### 2.4. TubeBot Motion Performance and Function Demonstration

Based on previous testing, the optimal TubeBot parameters were determined as follows: 50% mass fraction of NdFeB particles, 4 mm length, 45° initial magnetization angle, and a magnetization diameter of 1 mm. To further validate TubeBot’s performance in a vessel-like branched structure, experiments were conducted to observe TubeBot’s motion through channels of varying inner diameters, and its trajectory was recorded over time (Figure 4D and Video S3). Velocity profiles were calculated based on these trajectories (Figure S10). The results indicated that TubeBot’s speed increased with channel diameter in the four different inner diameter channels (0.8 mm, 1 mm, 1.2 mm, and 1.5 mm). TubeBot exhibited the highest speed in the 1.5 mm channel, while its speed was relatively lower in the 0.8 mm channel. Notably, TubeBot required more time to navigate branch points, largely due to manual adjustments of the rotating magnetic field to guide its path. Time-lapse images showed TubeBot’s trajectory at various time points, indicating stable performance even in channels with diameters close to TubeBot’s outer diameter. When changing the path, the TubeBot moves forward and backward by changing the rotation direction of the magnetic field, which shows that the TubeBot is highly maneuverable even in a variable-path channel.

TubeBot’s adaptability to varying channel diameters was further tested using a stepwise tapered channel model constructed from silicone tubes of different diameters (1 mm, 2 mm, 3 mm) (Figure S11 and Video S3). TubeBot successfully adjusted to changes in diameter and crossed intermediate steps regardless of whether it was transitioning from smaller to larger diameters or vice versa.

To evaluate TubeBot’s cargo-loading and releasing capability, TubeBot loaded with alginate hydrogel was used to demonstrate its movement through a maze phantom and release at the target point (**Figure 5A**(i)). Alginate loaded with fluorescent nanoparticles was used to visualize the process, and TubeBot successfully carried and maintained the integrity of the material (Figure 5A(ii)). Alginate, a hydrogel that can be compressed in aqueous solution after cross-linking. TubeBot’s crawling is primarily driven by body deformation, and the loaded alginate did not hinder TubeBot’s deformation under the magnetic field (Figure S12A). Crawling tests in a sloping tube further demonstrated TubeBot’s ability to move in complex environments while carrying the cargo (Figure S12B). In the maze phantom, TubeBot navigated through different pathways (with branch angles ***θ*1**, ***θ*2**, and ***θ*3**) and successfully reached the target (Figure 5B and Video S4). TubeBot efficiently navigated through branches of varying angles while loaded, demonstrating its load-carrying capacity and efficient movement in complex channels (Figure S13). After reaching the target point, TubeBot released the alginate in a 37.5°C environment over 24 hours, verifying its release capability (Figure 5C and Video S4). As time progressed, alginate was gradually released under enzymatic action, with the distribution of fluorescent nanoparticles at 85 minutes demonstrating the release process (Figure 5C(ii)). After 24 hours, the release of alginate had ended and the fluorescent nanoparticles were scattered around the target site (Figure 5C(iii)). This experiment confirmed TubeBot’s potential as a drug delivery platform.

**Figure 5.**
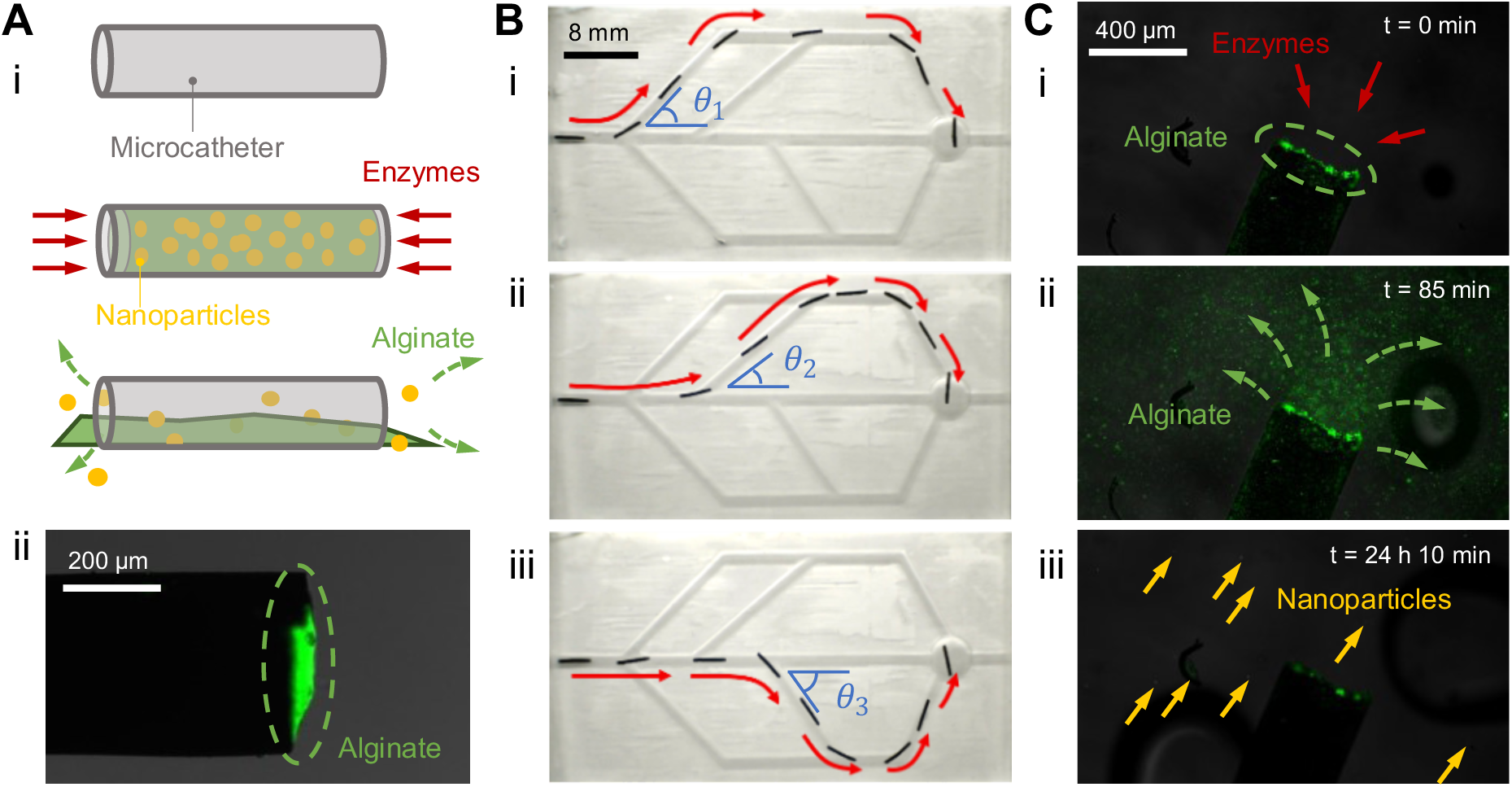
TubeBot function demonstration. (A) (i) Schematic of alginate and fluorescent particles loading and release by the TubeBot. (ii) Fluorescence image of TubeBot loaded with alginate. (B) TubeBot loaded with alginate navigating to the target point along different paths in the channel. (C) The TubeBot releases alginate at the target point over 24 hours. (i) *t* = 0 min, alginate is completely loaded into TubeBot. (ii) *t* = 85 min, alginate is degraded by the enzymes and releases fluorescent nanoparticles stably. (iii) *t* = 24 h 10 min, alginate has been degraded and the fluorescent nanoparticles are scattered around the target chamber.

To assess the effects of TubeBot’s crawling on cell activity, experiments were conducted to evaluate its movement over cell layer. TubeBot was tested within a bovine oviduct epithelial cells (BOECs) model (Figure S14). TubeBot crawled smoothly over the BOECs layer with continuous reciprocating movement, without causing any interference or adhesion to the cells. Fluorescence staining was used to evaluate cell viability, and images showed that the BOECs layer remained intact after multiple passes by TubeBot, with no evidence of widespread cell damage or significant reduction in cell viability. These findings indicate that TubeBot’s movement over the BOECs layer was gentle enough to avoid physical damage, validating its safety in biological environments. Details of the experiments are in the “TubeBot testing in BOECs model” section of the Supporting Information.

### 2.5. The Microcatheter Robot

To enhance the active propulsion capabilities of the microcatheter robot for navigation in complex tissue structures and effective connection with medical tubings, a design similar to TubeBot was developed, but with increased length. The microcatheter robot features a magnetization radius (r) of 0.5 mm, wrapped in three helical turns, with a total length of 15 mm (**Figure 6A**(i)). The innermost loop has a magnetization diameter of 1 mm, which gradually increases for the subsequent loops, but both ends of the magnetized catheter remain aligned on the same axis, with an initial magnetization angle set to 45°. The internal magnetic moment distribution after magnetization, shown in Figure 6A(ii), confirms the diameter variation.

**Figure 6.**
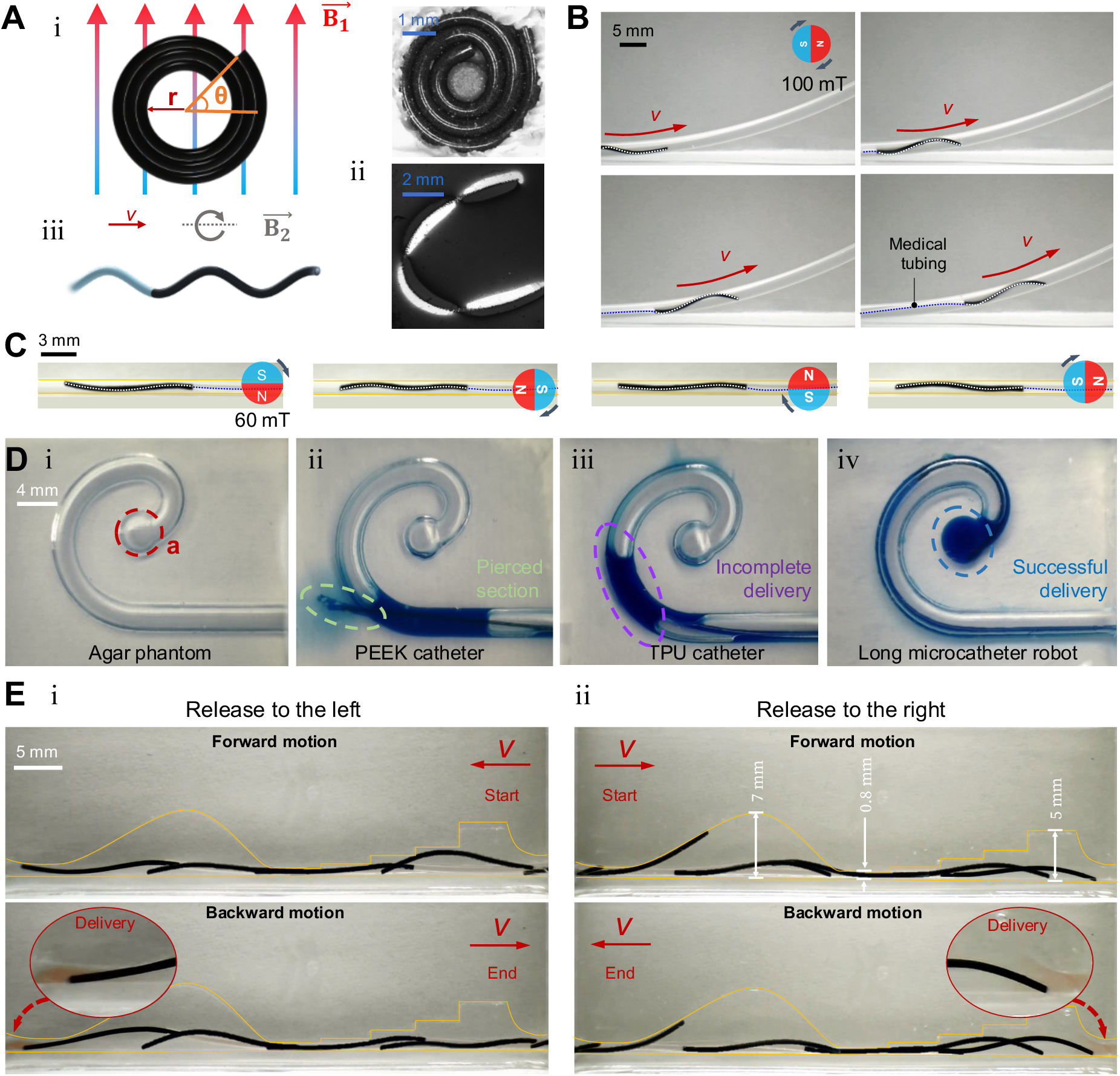
Microcatheter robot optimization and function demonstration. (A) (i) The microcatheter robot is fixed in a mold (*r* = 1 mm) and magnetized using a uniform magnetic field 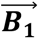. (ii) The visualization of the magnetization distribution by MOKE microscopy, (iii) The microcatheter robot connected to a medical tubing and is propelled forward by a rotating magnetic field 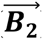. (B) The microcatheter robot crawls upward through a sloping silicone tube (*ID* = 2 mm) without external pulling. (C) Postures of the microcatheter robot at various angles of the rotating magnetic field in a silicone tube (*ID* = 1 mm). (D) Advancement of different medical tubes in an agar phantom. (i) Overview of the complete agar phantom. (ii) The PEEK medical tubing (*d* = 0.38 mm) pierces the agar phantom channel. (iii) The TUP catheter (*d* = 0.64 mm) is stuck in the middle of the channel. (iv)The microcatheter robot successfully crawls to target point A and transfers liquid. (E) The microcatheter robot demonstrates robustness by crawling and pushing forward in a complex channel. The yellow line outlines the channel. (i) The microcatheter robot crawls forward from right to left, then backward from left to right, effectively transporting liquid (red). (ii) The microcatheter robot connected to medical tubing, crawls forward from left to right and backward from right to left within a complex channel.

Upon connection to medical tubing, the microcatheter robot achieved terminal active propulsion under a rotating magnetic field, propelling the medical tubing forward (Figure 6A(iii)). The microcatheter robot was placed in an inclined channel with an inner diameter of 2 mm and driven by a rotating magnetic field of 100 mT. The results demonstrated that the microcatheter robot was able to overcome gravity and successfully dragged the medical tubing upward without external assistance (Figure 6B), confirming its capability of terminal active propulsion. The microcatheter robot was subjected to different magnetic fields in a channel with a diameter of 1 mm, resulting in deformation over a magnetic field rotation cycle (Figure 6C). The white dashed line indicates the posture of the magnetic catheter component of the microcatheter robot, which took on a complete sinusoidal shape, generating continuous propulsive forces. The blue dashed line indicates the posture of the medical tubing. The ability of the microcatheter robot to deliver material in simulated biological environments was assessed using an agar phantom, which shares similar characteristics with human soft tissue.^[26]^ The performance of different catheters in delivering to target point a was compared (Figure 6D). The polyetheretherketone (PEEK) catheter failed to navigate through the agar phantom as its stiffness prevented it from bending to follow the path, resulting in puncture (Figure 6D(ii)). The thermoplastic polyurethane (TPU) catheter failed to reach the target point and got stuck in the middle of the phantom because, without terminal deflection, the static friction between the catheter and the phantom impeded its advancement (Figure 6D(iii)). In contrast, the microcatheter robot demonstrated superior navigation capabilities in the agar phantom, effectively crawling to target point a and successfully delivering fluid without damaging the phantom (Figure 6D(iv)). These findings highlight the microcatheter robot’s potential for safe material delivery in soft tissue environments, with performance significantly surpassing that of traditional catheters.

To evaluate the robustness of the microcatheter robot’s motion capabilities, it was placed in a complex channel to perform both forward and reverse crawling to deliver fluids (Figure 6E and Video S5). The channel outline was highlighted with yellow lines, and the upper portion of the channel featured an asymmetric structure, as detailed in Figure S15. During the leftward delivery experiment (Figure 6E(i)), the microcatheter robot crawled forward from right to left, passing through a descending step and an arched section, with the narrowest part having a channel height of 0.8 mm. During the return process, the microcatheter robot successfully delivered red fluid along the edge of the channel, before moving in reverse from left to right. In the rightward delivery experiment (Figure 6E(ii)), the microcatheter robot navigated through the arched section first, followed by the step section, delivering fluid and then reversing from right to left. The experimental results indicate that regardless of the direction of movement in the complex channel, the microcatheter robot was able to adapt to changes in the channel structure and achieve effective fluid delivery. The robot’s robustness in navigating complex channels demonstrates its high adaptability and delivery stability, further highlighting its potential advantages for practical clinical applications.

### 2.6. Microcatheter Robot Navigation Propulsion Demonstration in Phantoms

In traditional procedures, navigating to deep regions of the bronchial tree often involves multiple complex steps. Typically, this process uses a flexible bronchoscope that is inserted into the trachea and gradually advanced to the deeper branches of the bronchial tree.^[27]^ The flexible bronchoscope is widely used to access smaller airways for visual inspection, sample collection, or drug delivery. Throughout the procedure, a guiding catheter often serves as an anchor for deploying and stabilizing other catheters or tools, with the flexible catheter advancing toward the target area.^[28]^ Given the complexity of the bronchial tree, additional guidewires or instruments may be needed, increasing the difficulty of the operation and the strain on the patient’s airways. This process demands considerable skill and often requires multiple adjustments, particularly when anatomical abnormalities or narrow airways are present. Multiple insertions and equipment changes within the bronchial tree not only prolong the procedure but also increase the risk of damage to the bronchial mucosa, bleeding, and other complications.^[29]^ Minimizing the use of catheters and guidewires can enhance surgical efficiency and safety, reducing potential risks and overall procedural time.

Here, we evaluated the microcatheter in various 3D models to demonstrate its capability as an integrated device for navigating complex anatomical structures (**Figure 7** and Video S6). In the initial demonstration, the microcatheter navigated through a simplified 3D channel with multiple side branches, as shown in the side view (i) and top view (ii) of Figure 7A. The microcatheter entered the main channel at *t* = 0 s, advanced into channel 1 (*d* = 1.5 mm), and traversed bends ranging from 10-40° in 3D space within 100 s (operated by a user with limited experience). This highlights the device’s ability to navigate complex branching pathways within a certain timeframe, facilitating further clinical procedures.

**Figure 7.**
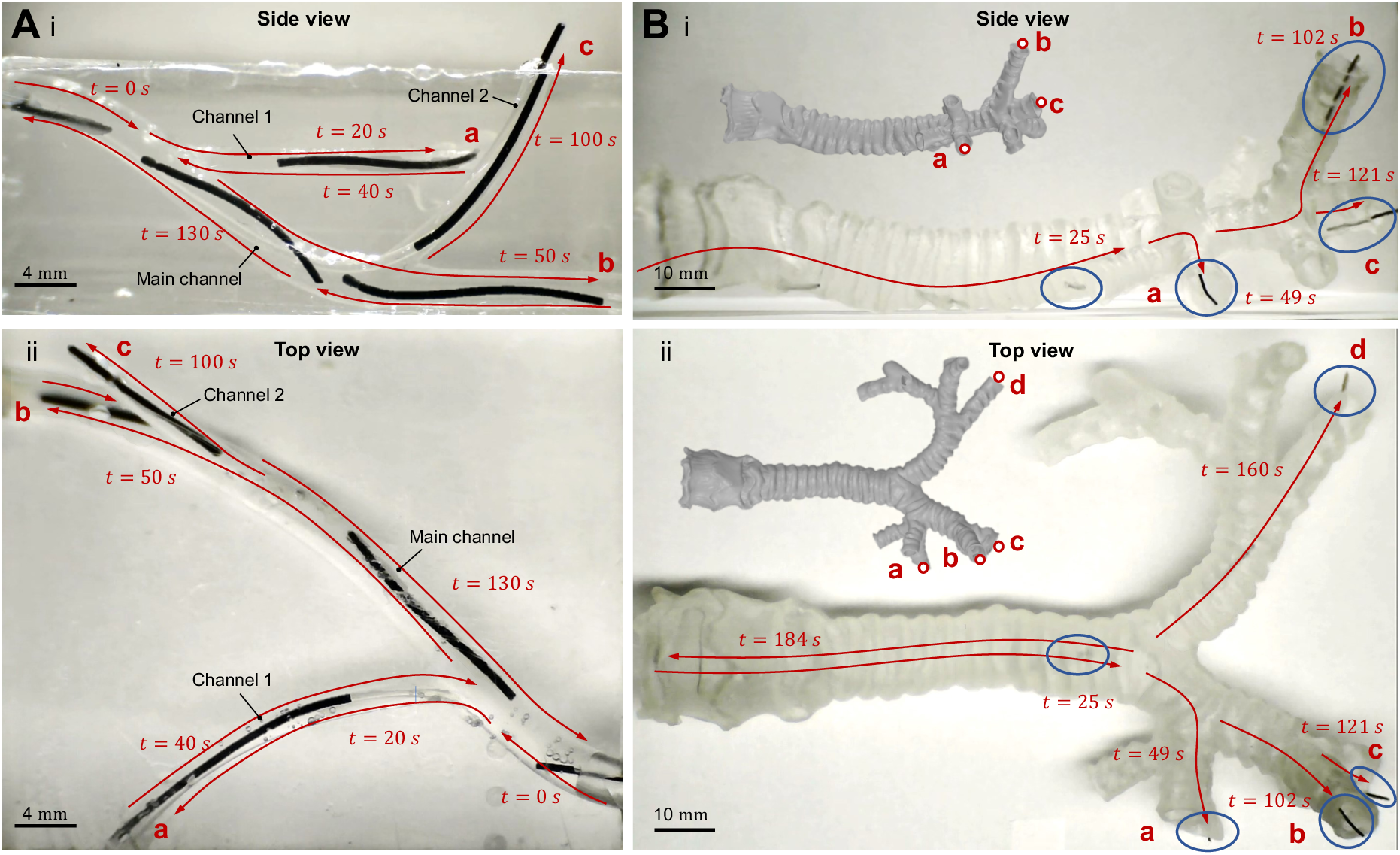
Demonstration of the microcatheter robot platform navigation through the 3D phantoms. (A) Navigation and crawling of microcatheter robot in a 3D channel, shown in side view (i) and top view (ii). The microcatheter robot enters the main channel (*d* = 3 mm) at *t* = 0 s, driven by a rotating magnetic field. At *t* = 20 s, the field is adjusted to guide it into side channel 1 (*d* = 1.5 mm) towards target point a. By *t* = 40 s, the microcatheter robot exits side channel 1, and at *t* = 50 s, it is guided to target point b in the main channel. At *t* = 100 s, the microcatheter robot enters side channel 2 (*d* = 1.5 mm) and reaches target point c. Finally, at *t* = 130 s, the microcatheter robot exits the main channel through a combination of the rotating magnetic field and manual pulling. (B) Navigation and crawling of the microcatheter robot in a full-scale 3D bronchial tree phantom, shown in (i) side view and (ii) top view. The microcatheter robot reaches the bronchial site at *t* = 25 s, enters the left bronchial tree at *t* = 49 s, and reaches target point a. By *t* = 102 s, it advances to target point b, and by *t* = 121 s, it reaches target point c. At *t* = 160 s, the microcatheter robot exits the left bronchial tree, enters the right bronchial tree, and reaches target point d. Finally, the microcatheter robot exits the bronchial tree at *t* = 184 s, driven by the rotating magnetic field and manual pulling.

Moreover, a comparison of the microcatheter robot’s performance in side branches using manual pushing and terminal propulsion methods (Figure S16) showed that manual pushing combined with magnetic steering could lead to jamming in narrow or sharply curved channels. In contrast, terminal crawling under magnetic navigation provided robustness, allowing the microcatheter robot to traverse challenging sections successfully.

Further tests in a complete 3D bronchial tree model assessed the microcatheter’s ability to follow anatomically accurate and complex paths (Figure 7B). Both the side view (i) and top view (ii) demonstrate the robust movement of the microcatheter through the bronchial model. After entering the main bronchus, the robot crawled through bends ranging from 10-60° and covered a distance of over 316 mm within 160 seconds. These demonstrations underscore the microcatheter’s ability to navigate challenging environments, maintain stability, and achieve directional control—all of which are crucial for minimally invasive applications in medical settings and lay the groundwork for in vivo experiments.

### 2.7. Ex Vivo and In Vivo Propulsion of Microcatheter under US Imaging Guidance

To enable real-time visualization of the microcatheter robot, an ultrasound (US) imaging system was used to monitor its movement within an ex vivo phantom model and ex vivo mouse uterine tissue, thereby evaluating its navigation capability and stability in complex biological environments **(Figure 8** and Video S7). Figure 8A shows the microcatheter robot’s movement in the tubing model. As depicted in Figure 8A(i), the experimental setup included a rotating magnetic field generated by a permanent magnet coupled to a step motor to drive the robot’s motion, with an ultrasound probe in contact with the tube using ultrasound gel to ensure signal transmission. The actual settings are shown in Figure S17. The US imaging (Figure 8A(ii)) captured the microcatheter robot (highlighted in yellow) as it dragged a medical tubing (highlighted in brown) under magnetic actuation. Speed data presented in Figure 8A(iii) demonstrated consistent movement speeds across different microcatheters under identical magnetic field conditions, indicating that the microcatheter robot maintained stable crawling velocity and effectively completed the task of crawling and dragging the medical tubing within the channel phantom setup.

**Figure 8.**
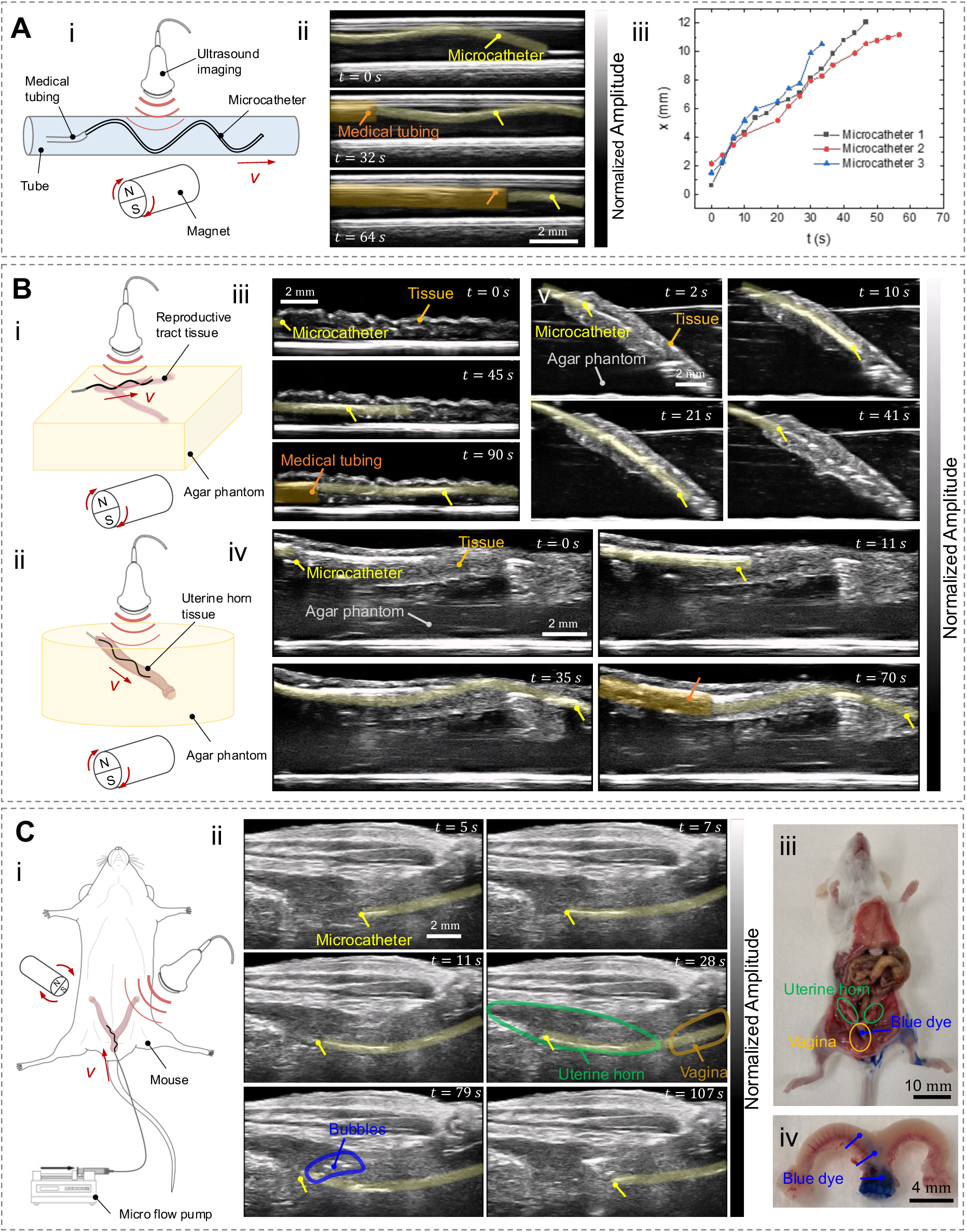
Ex vivo and in vivo actuation and imaging of the microcatheter robot. (A) (i) Schematic of the tube phantom setup. (ii) US image showing the microcatheter robot (yellow transparent is the microcatheter robot, brown transparent is medical tubing) crawling through the tube phantom while pulling a medical tubing. (iii) Speed of the microcatheter robot within the tube phantom under a 100 mT magnetic field at 7 Hz. (B) US imaging of the microcatheter robot in uterine tissue ex vivo. (i) Schematic showing uterine tissue positioned on the surface (reproductive tract tissue) and (ii) within (uterine horn tissue) an agar phantom. (iii) US imaging of the microcatheter robot crawling through the uterus on an acrylic surface. (iv) US imaging of the microcatheter robot crawling through the uterus on the agar phantom, maintaining its 3D structure. (v) US imaging of the microcatheter robot crawling inside the uterus embedded in the agar phantom, reaching the bottom under US guidance. (C) Schematic of in vivo actuation in a mouse uterus under US guidance. (i) US imaging of the microcatheter robot crawling in the uterus and delivering dye. (ii) The microcatheter robot reaches the uterus via the vagina and delivers dye through medical tubing. (iii) Post-mortem examination showing uterine staining. (iv) Blue dye marks the site of delivery on the left side of the uterus.

Further experiments evaluated the microcatheter robot’s movement within ex-vivo mouse uterine tissue (Figure 8B). The experimental setups involving uterine tissue placed on an agar surface and embedded in agar are shown in Figs. 8B(i) and 8B(ii). Figure 8B(iii) displays US imaging of a microcatheter robot crawling within mouse uterine tissue on an acrylic surface, where significant signal reflection from the bottom interferes with the observation of the microcatheter robot. The use of agar effectively reduced signal reflection from the acrylic base, resulting in clearer US imaging of the microcatheter robot (Figure 8B(iv)). Figure 8B(v) further demonstrates the robot’s movement within uterine tissue embedded in an agar phantom, and agar has acoustic properties similar to those of biological soft tissue.^[26]^ The preparation of uterine horn tissue embedded in agar allowed the formation of an environment resembling a three-dimensional in vivo structure after agar solidification. Under US imaging guidance, the robot successfully crawled along the uterine horn, reached the base of the phantom, and ultimately crawled backward to exit the phantom without any significant movement hindrance. These experimental results validated that under real-time US imaging guidance, the microcatheter robot could reach target points within both the channel and ex-vivo biological tissue, exhibiting consistent speed and stability across different environments.

To demonstrate the clinical applicability of this technology, an in vivo feasibility study was conducted on a mouse model. The female mouse reproductive system is often used as a preliminary model for the human reproductive system, despite differences in size and anatomy, as their basic functions are similar.^[30]^ The microcatheter robot was employed to navigate and deliver fluid within the mouse reproductive tract, with its movement and positioning monitored in real-time via US imaging (Figure 8C). The experimental setup (Figure 8C(i)) involved the microcatheter robot being inserted through the vagina and gradually navigating and clawing to the uterine horn under the auction of a rotating magnetic field. The microcatheter robot is connected to medical tubing and a syringe containing blue dye (Figure S18). Once positioned, fluid delivery to the target point was facilitated by a microfluidic pump. US imaging illustrated the robot’s movement within the mouse reproductive system (Figure 8C(ii)). At *t* = 5 s, the robot entered the mouse’s vagina (yellow-marked region), and by *t* = 28 s, it had successfully navigated to the uterine horn (green-marked region). The US images demonstrated the robot’s movement from the vagina through to the uterine horn. At *t* = 79 s, blue dye was pumped through the microcatheter robot’s tip and released into the uterine horn. The movement and dispersion of white bubbles within the dye (blue-marked region) indicated smooth, unobstructed dye delivery. After the experiment, euthanasia was performed, and dissection confirmed the successful delivery of the dye to the uterine horn (Figure 8C(iii) and 8C(iv)). The results demonstrated that the microcatheter robot could effectively navigate and deliver fluids within a complex in vivo reproductive environment, opening new ways for minimally invasive drug delivery or more effective in vivo assisted insemination.

## 3. Conclusion and Outlook

This research proposes a high-yield fabrication method for multifunctional magnetic microcatheter robots, offering rapid and cost-effective customization for various biomedical applications. The platform’s programmable magnetization enables adaptability across different robotic configurations, enhancing navigation and propulsion in complex anatomical structures. Unlike traditional rigid guidewires requiring frequent adjustments, our platform utilizes soft-material-based magnetic microcatheters for remote magnetic navigation with embedded wires to tune stiffness, reducing tissue trauma risk. The wave crawling propulsion mechanism increases robustness and enables navigation under standard ultrasound imaging. Compared to existing end-propelled catheters, our microcatheter robots offer advantages in size, manufacturing process, and propulsion performance while maintaining functional working channels at the micrometer scale (Table S2).

The crawling mechanism effectively converts static friction to dynamic friction when the microcatheter becomes stationary or obstructed. Through sinusoidal deformation, each contact point pulls the microcatheter robot in reverse, helping it navigates bifurcations or narrow passages. When stuck at sharp turns, the global rotating magnetic field induces sinusoidal deformation, increasing tissue contact points while decreasing overall contact area. In this case, dynamic friction becomes less than static friction, enabling forward crawling and assisted propulsion, allowing the robot to extricate itself (Figure S16).

Compared to continuum robots driven by axial force transmission, our distal propulsion approach mitigates perforation risk since proximal forces don’t directly affect the tip. When the robot becomes stuck, external force converts to wave deformation rather than transmitting axial force to the tip. Wave crawling transforms elastic deformation energy into frictional force through point contact, propelling the microcatheter forward. The soft-tipped design further reduces perforation risk.

Although wave crawling induces slight vibrations, the microcatheter’s soft nature minimizes tissue damage risk, as demonstrated in tests on BOEC layers (Figure S14). Further studies are needed to assess vibration impacts in different tissue environments. TubeBot observations revealed that propulsion combines sinusoidal undulation and lateral sway, which could enhance adaptability in future designs.

Currently, except for the wireless TubeBot, all microcatheters require manual propulsion, though the platform can accommodate commercial electric actuators for automated propulsion. Effective wave crawling in larger luminal structures remains challenging due to insufficient tissue contact. However, static magnetic fields with manual pushing can still facilitate navigation, while dynamic fields with increased oscillation amplitudes help navigate complex anatomical regions (Figure 7B).

While surface coatings weren’t explored in this work, future research should consider biocompatibility enhancement and friction optimization, especially for long-duration surgeries. Conventional designs using rigid distal magnets require a trade-off between flexibility and control. By embedding magnetic particles in the polymer, our platform achieves enhanced flexibility at the cost of a lower magnetic moment, necessitating stronger driving fields. The magnetic navigation system of the current platform primarily relies on permanent magnets to provide the field strength required for effective operation (100 mT), allowing efficient functioning. For clinical applications, achieving the desired magnetic field strength at the in vivo device location requires stronger permanent magnets, potentially assisted by robotic arms to follow the device.^[31]^ This increases the working space of the system and may also lead to interference with other surgical room equipment. Future clinical applications may benefit from electromagnetic approaches, which would allow for more precise and adjustable control, although implementing such a system requires careful consideration of space, interference with other equipment, and overall practicality.^[13a]^

Compared to traditional systems, the lack of haptic feedback remains a challenge. Surgeons are accustomed to the tactile feedback provided by conventional guidewires, which assists in navigating anatomical paths and reducing tissue damage. Although some devices now integrate pressure sensors to provide feedback, the forces generated by flexible microcatheters are usually too small, and in the complex channel, the pressure is dissipated to the outside through deformation and friction sensing.^[32]^ Therefore, it cannot provide useful data during surgery. In the future, contact stress should be calculated based on US images or other real-time imaging, and force sensors should be integrated into the catheter tip to improve navigation accuracy. To enhance catheter tracking accuracy in vivo, on-board electronic sensors are integrated for magnetic tracking.^[33]^ The incorporation of various sensors can improve the catheter’s safety and maneuverability, but it also presents challenges related to catheter size and internal lumen space. A key advantage of the proposed microcatheter platform lies in its capability for rapid, cost-effective production across multiple scales. By modifying the diameter of the tungsten wire used in manufacturing, microcatheters with varying sizes and working channel diameters can be produced to accommodate different spatial requirements, thereby enhancing adaptability for diverse biomedical applications.

Moreover, TubeBot demonstrated the ability to deliver drugs to confined anatomical regions (Figure 5), although challenges remain for long-distance transport. Lukas Hertle et al. developed a customizable, magnetic soft microrobot device using a microfluidic bionic extrusion approach. Upon reaching the target site, the device decomposes into a microrobot swarm to facilitate precise therapeutic delivery.^[34]^ This system illustrates the potential for flexible continuous robots and microrobots to operate cooperatively on a shared platform, enhancing each other’s functional efficiency. TubeBot and the microcatheter robot are both components of the microcatheter robot platform, with TubeBot seamlessly integrable with the microcatheter. In future applications, this integration promises a more efficient and targeted delivery mechanism, enabling precise, multi-scale interventions.

While these findings are promising, comprehensive in vivo testing and large-scale trials are necessary for regulatory approval and clinical deployment, requiring extensive risk assessment and validation to ensure safety and efficacy.

## 4. Experimental Section

### Preparation of Magnetic Microcatheter

NdFeB microparticles (MQFP-B-10215-089, Magnequench) with an average particle diameter of 5 μm were uniformly mixed into the PDMS resin base (Sylgard 184, Dow Corning) to prepare a ferromagnetic composite resin. NdFeB microparticles (10-50 weight % (wt %)) were mixed into the PDMS silicone base using a paddle stirrer (AR-100, Thinky) at 2000 rpm and stirred for 2 hours. After the resin solution mixed with NdFeB microparticles was cooled to room temperature, the curing agent was added in a weight ratio of silicone base to curing agent of 10:1 and mixed thoroughly. Next, the resin was degassed under vacuum to remove bubbles inside the resin. The magnetic microcatheter was prepared by Joule heating of a tungsten wire, and the preparation device is shown in Figure S1.^[35]^ The ferromagnetic composite resin was slowly transferred into the container, and a tungsten wire (7440-33-7, Goodfellow) with a diameter of 0.15 mm was passed through the container and fixed at both ends of the electric displacement platform. A constant current of 2.3 A was applied to the tungsten wire, and at the same time, the electric displacement platform (X-LSM200A-E03, Zaber) was used to move the tungsten wire upward at a constant speed. The Joule heat generated by the tungsten wire solidified the ferromagnetic composite resin on the tungsten wire. After the tungsten wire was removed, the microcatheter on the tungsten wire was manually and mechanically peeled off.

### Fabrication of Phantoms

The models were prepared using two different techniques. The first method was to print different resins using a 3D stereolithography printer (Mars 4, ELEGOO) to obtain a single flat or 3D model. The channels and molds were drawn using CAD software and 3D printed using resin (Build, Siraya Tech) and transparent resin (ABS-like Resin, ELEGOO). The complex channels on the flat surface were combined with a cover plate of Polymethyl Methacrylate (PMMA). The molds of 3D structures were molded using PDMS (10:1 mass ratio) to prepare different types of channels and bonded to glass slides to obtain ideal closed phantoms. The bonding process was to plasma treat the PDMS model and a clean glass slide for 1 min at a power of 99 watts, then adhere the PDMS model to the glass slide and bake it on a hot plate at 120°C for 20 min. In addition, the tracheal tree was 3D printed in one go using transparent resin. For the agar phantom, agar (9002-18-0, Sigma-Aldrich) and deionized water were mixed at a mass ratio of 3%, heated to 100°C, and manually stirred to dissolve. Wait until the agar solution is clear, cool to 40°C, and pour into the 3D printed mold. When cooled to room temperature, remove the agar phantom. The second method is to prepare the 3D channel phantom using the template method. The mold model was designed using CAD software, PMMA was cut accordingly to obtain mold parts, and finally hot glue (Bigstren) was used to get the mold. Put silicone tubes (Adtech polymer engineering Ltd) with different outer diameters (1.5 mm, 3 mm) into the mold, and use super glue (Stanger) to connect the silicone tubes. After standing for 1 hour, slowly pour PDMS (10:1 mass ratio) into the mold and vacuum exhaust. Place the mold containing PDMS on a hot plate at 65°C overnight to ensure that the PDMS is completely polymerized. After removing the glue between the mold parts, disassemble the mold to expose the PDMS model containing the silicone tube inside. Because the contact area between the silicone tubes is small and the silicone tubes are elastic, the silicone tubes inside the PDMS can be manually pulled out, and it is ensured that there is no solidified super glue blocking the internal channel.

### Magnetic Characterization

The magnetic moment density of ferromagnetic composite microcatheters with different NdFeB particle concentrations was measured using a vibrating sample magnetometer (DMS 1660, ADE Technologies). The samples were prepared by cutting microcatheters with an outer diameter of 300 μm into 5 mm lengths and fixing them into the sample holder of the magnetometer. The external magnetic field was applied up to 3 T and cycled twice. When the applied external magnetic field was zero, the residual magnetic moment of the sample was measured and then divided by the sample volume to obtain the magnetization or magnetic moment density of the sample.

### Mechanical Testing

The dog-bone-shaped ferromagnetic composites with different NdFeB particle concentrations were prepared by 3D printing molds. The specimens used for tensile testing had a test width of 4 mm and a gauge length of 17 mm. The specimens were tensile tested on a mechanical testing machine (Z005, Zwick/Roell) using a 2.5 KN load cell at a test rate of 50 mm/min.

### Radiography Characterization

To assess the distribution of the NdFeB magnetic particles in the microcatheter X-Ray computed tomography (XCT) (Phoenix nanotom m, General Electric) has been conducted. Scans were performed at 80 kV and 120 µA with a voxel size of 1 µm. Samples were prepared by cutting microcatheters with 300 μm outer diameter into roughly 5 mm lengths and clueing them on sample holder for XCT. To illustrate the tomographic reconstruction in different magnifications and views, including longitudinal, oblique, cross-sectional, and local magnified view, the Software VGStudioMax 2023.4 has been used.

### Magnetization of Microcatheter Robots

After the microcatheter embedded with NdFeB particles is peeled off from the tungsten wire, the microcatheter is personalized for different functional applications. All the demonstrated magnetic microcatheter prototypes are uniformly magnetized by a pulsed magnetic field (3T) generated by a pulsed magnetizer (IM2525MAC, Magnet-Physik) so that the NdFeB particles are magnetically saturated. The guiding microcatheter is magnetized axially, and the microcatheter is fixed in the vertical groove of the 3D-printed mold during magnetization. The magnetization of the TubeBot adopts a programmed magnetization curve that approximates a sinusoidal magnetization curve. The microcatheter is fixed to the mold at 45° with the starting end and the magnetic field direction, close to a cylinder with a diameter of 1mm, and ensures that the entire microcatheter is in one plane. The magnetization of the microcatheter robot is similar to that of TubeBot. The starting end of the microcatheter robot is an axial magnetized end with a length of 1 mm as the navigation end, and then the remaining microcatheter part is programmed to be magnetized using a sinusoidal magnetization curve. It is also necessary to ensure that the entire microcatheter is in one plane during magnetization.

### Magneto-optical Kerr effect Characterization

A magneto-optical indicator film with perpendicular anisotropy (PMOIF) is employed to visualize the magnetization distribution within a microcatheter. The core of this PMOIF is a transparent magnetic garnet film, which is deposited on a transparent gadolinium-gallium-garnet (GGG) substrate and coated with a mirror layer (Fig. S6). In the absence of an external field, the magnetic garnet film naturally forms a ground state of alternating, up-and-down magnetized band domains. When positioned on the surface of a magnetic sample, the stray field originating from the sample’s poles causes modifications in the perpendicular garnet band domain, changing the “density” of up- or -down-domains. By using a magneto-optical wide-field Kerr microscope configured for polar sensitivity under low-resolution settings—where the individual garnet domains are not visible—a polar Faraday contrast is produced in the reflection polarization microscope.^[36]^ This contrast pattern mirrors the pole arrangement within the cylindrical microcatheter revealing the magnetization distribution in it.^[37]^

### Microcatheter Robot’s Cargo Loading and Release

A microcatheter with a length of 4 mm was programmed to magnetize with a sinusoidal magnetization curve, and the microcatheter was treated with plasma (99 watts) for 1 min. Alginate (9005-38-3, Sigma-Aldrich) was mixed into deionized (DI) water at a mass ratio of 5:100 to prepare an alginate solution. The alginate solution was diluted to 0.5% (mass ratio) and vortex-mixed with fluorescent nanoparticles (26988, Duke Scientific Corporation) at a volume ratio of 30:1. One end of the microcatheter was connected to a capillary (Hilgenberg GmbH) with a tip outer diameter of approximately 140 μm, and the alginate solution containing fluorescent nanoparticles (particle size of 200 nm) was injected into the microcatheter through a microinjection pump (Cetoni GmbH). The microcatheter loaded with alginate solution was placed in a CaCl_2_ (10043-52-4, Sigma-Aldrich) solution with a concentration of 200 mM for 30 seconds, and then taken out and placed in DI water for storage. The release of cargo from the Microcatheter robot was achieved by the degradation of alginate. The enzyme (Alginate Lyase, 9024-15-1, Sigma-Aldrich) was prepared with 1X phosphate buffered saline (PBS) at a mass ratio of 1%. The microcatheter robot was placed in an environment of 37°C and the enzymatic degradation solution was added to make the enzyme concentration (10^%&^ mg/ml) stable in the overall environment. Immediately, the sample was placed under a fluorescence microscope and photographed, mainly using Fluorescein Isothiocyanate (FITC) and Bright filter, and photographed every 5 minutes for 24 hours.

### Microcatheter Connects with Medical Tubing

One end of the microcatheter was coated with a biocompatible UV-curable adhesive (Loctite AA 3942, Henkel), connected to one end of the medical tubing (*ID* = 0.38 mm, polyurethane medical tubing, Nordson Medical), and irradiated with a UV lamp for 1 min at a wavelength of 365 nm and a power of 100 mW/cm^’^.

### Magnetic Actuation and Demonstration

In the deflection demonstration experiment for the guiding microcatheter, a uniform magnetic field was applied using a pair of Helmholtz coils. The guiding microcatheter was positioned vertically at the center of the coils, where it initially drooped due to gravity. Upon applying the magnetic field, magnetic torque induced deflection in the microcatheter, and by modulating the local stiffness of the guiding microcatheter, different spatial deflection ranges were achieved.

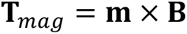

Here ***T_mag_*** is the magnetic torque, **m** is the magnetic dipole moment, **B** is the strength of the external magnetic field generated by the coils. Copper wires of different lengths were inserted into the guiding microcatheter, and the position of the tip of the guiding microcatheter under different magnetic field strengths was recorded.

For the navigation and crawling demonstrations proposed in this article, cylindrical NdFeB magnets (STM-10x50-N-D, magnets4you GmbH) with a diameter of 10 mm and a length of 50 mm were used to apply the magnetic field required for long-distance crawling. For magnetic steering and navigation, the magnets were manually manipulated to change their position and direction, and the magnets were kept at the proximal end of the guiding microcatheter when the guiding microcatheter was advanced, thereby changing the direction and strength of the applied magnetic field. For crawling, the position and direction of the magnet need to be manually manipulated, and the magnet speed can be changed by changing the parameters of the stepper motor driver controller (GY20512-FBA, PEMENOL). When the microcatheter robot is connected to the medical tube for crawling, the magnet needs to be manually manipulated to keep it at the proximal end of the microcatheter robot, and the entire body is advanced by pushing the distal end combined with the crawling of the proximal end, and the steering and navigation tasks are controlled by visual feedback.

### Ex Vivo Experiments

The mouse uteri used in this article were obtained from mice sacrificed by the Transgenic Core Facility of the Max Planck Institute of Molecular Cell Biology and Genetics (Dresden) under license numbers TVV 5712021 (Etablierung einer neuartigen, nicht-invasiven Methods für den Embryotransfer mit innovativen Mikrorobotik-Tools). The mouse uteri used for ultrasound imaging (US imaging) were removed after mouse sacrifice, placed in PBS, stored at 5°C, and used within 24 hours. In the first ex vivoexperiment, the mouse uterus was placed directly on the surface of acrylic support covered with ultrasound gel (Aquasonic) for signal transduction and directly subjected to ultrasound imaging. In the second protocol, the mouse uterus was placed on the surface of agar and covered with ultrasound gel on top. In the last protocol, the mouse uterus was placed inside agar, covered with ultrasound gel on the surface of agar, and subjected to ultrasound imaging. 100 ml of DI water was heated to 80°C, and then 3 g of agarose was slowly added and gently mixed to form a homogenous solution (3%wt). A microtube with an outer diameter of about 400 μm was inserted into the mouse uterus for support. After it was placed in a container, an agarose solution at about 40°C was poured in. After cooling to room temperature and becoming a gel, the microtube was taken out. The magnetic drive of the microcatheter robot was achieved using a magnet placed under the phantom, with a rotation frequency of 7 Hz and a magnetic field strength of about 100 mT. At the same time, to visualize such microcatheter robots within the tissue using existing imaging technology, a US imaging system (FUJIFILM VisualSonics) was used in the experiment. The experiment was conducted using a 256-element transducer with a center frequency of 25 to 57 MHz, and the captured data was collected and reconstructed using onboard post-processing.

### In Vivo Experiments

The mice used in this article were obtained from the Transgenic Core Facility of the Max Planck Institute of Molecular Cell Biology and Genetics (Dresden) under licenses TVV 5712021 (Etablierung einer neuartigen, nicht-invasiven Methoden für den Embryotransfer mit innovativen Mikrorobotik-Tools). They comply with the guidelines of the Federation of European Laboratory Animal Science Associations (FELASA) and the National Law on Laboratory Animal Experiments (Law No. 18.611). Twelve-week-old mice were used for in-utero imaging. All mice were anesthetized with 1.5-2% isoflurane. O_’_ flow (maintained at 1 − 2 ml/min) was mixed with isoflurane. To optimize the coupling of ultrasound gel, the hair on the abdomen of the mice was removed using a commercially available depilatory cream. Ultrasound imaging was also performed using a US imaging system. The experiment was conducted using a 256-element transducer with a center frequency of 25 to 57 MHz. After finding the mouse uterus in the ultrasound imaging system, the robot entered the mouse body through the vagina under the drive and navigation of the magnet. When the tip of the long microcatheter robot reached the bottom of the uterus, the crawling drive was stopped, and a microfluidic pump (Cetoni GmbH) was used to deliver blue dye (6104-58-1, Sigma-Aldrich) to the inside of the uterus through the medical tubing to the long microcatheter robot. The magnet was reversely controlled to rotate and drive the long microcatheter robot to exit the uterus.

## Supporting Information

Supporting information is available from the Wiley Online Library or from the author.

## Acknowledgements

We gratefully acknowledge the contributions of the following individuals and teams: Thomas George Woodcock (IFW-Dresden) for assistance with sample magnetization; Volker Neu and Nicolas Perez Rodriguez (IFW-Dresden) for their support in magnetic characterization; and Nicole Geißler (IFW-Dresden) for conducting tensile testing of the samples. We also thank the cleanroom team at IFW-Dresden for their assistance in microscopic characterization and 3D printing of molds. Additionally, we acknowledge Carla Ribeiro (IFW-Dresden) for providing bovine oviduct epithelial cells and Ronald Naumann (MPI-CBG) for supplying murine ex vivo uterine tissues; and the team of experimental animal platform (DZNE) for their support in animal experiments. The authors gratefully acknowledge the financial support received from the European Union’s Horizon 2020 research and innovation program (ERC Starting Grant Nr. 853609). And the HORIZON-MSCA-2022-COFUND-101126600-SmartBRAIN3. L.B. is co-financed with tax funds based on the budget adopted by the Saxon state parliament under SAB-Nr.: 100632843

## Conflict of Interest

Zhi Chen, Boris Rivkin, and Mariana Medina Sánchez have filed a provisional patent application on the basic principle and design of the microcatheter preparation device. The other authors declare that they have no competing financial interests.

## Data Availability Statement

The data that support the findings of this study are available from the corresponding author upon reasonable request.

## Funding

The authors gratefully acknowledge the financial support received from the European Union’s Horizon 2020 research and innovation program (ERC Starting Grant Nr. 853609). And the HORIZON-MSCA-2022-COFUND-101126600-SmartBRAIN3. L.B. is co-financed with tax funds based on the budget adopted by the Saxon state parliament under SAB-Nr.: 100632843.

## Notes

### Competing Interest Statement

The authors have declared no competing interest.

