## Supplementary material for "Self-Propelling Adaptive Robotic Microcatheters Enabled by Scalable Fabrication for Intracorporeal Navigation": Supporting Information.pdf

**List of acronyms and abbreviations**

| | |
|---|---|
| 3D | Three-dimensional |
| MIS | Minimally invasive surgery |
| HSG | Hysterosalpingography |
| RMN | Remote magnetic navigation |
| mCR | Magnetically driven continuum robot |
| mSCR | Magnetic soft continuum robot |
| TubeBot | The tubular microrobot |
| PDMS | Polydimethylsiloxane |
| NdFeB | Neodymium-iron-boron |
| SEM | Scanning electron microscopy |
| VSM | Vibrating sample magnetometry |
| MOKE | Magneto-optical Kerr effect |
| PBS | Phosphate-buffered saline |
| BOECs | Bovine oviduct epithelial cells |
| PEEK | Polyetheretherketone |
| TPU | Thermoplastic polyurethane |
| US | Ultrasound |
| MicroCT | Micro Computed Tomography |

**Supplementary Note:**

**Landau-Levich-Derjaguin theory**

According to the Landau-Levich-Derjaguin theory,[1]

$$\mathrm{h} = \frac{1.34rCa^{\frac{2}{3}}}{1 - 1.34Ca^{\frac{2}{3}}}$$

where h is the thickness of the resin mixture coating, which corresponds to the wall thickness of the microcatheter after curing. r is the radius of the tungsten wire, Ca is the capillary number,

$$\mathrm{Ca} = \frac{\mu V}{\sigma}$$

with μ as the dynamic viscosity of the resin mixture, V the pulling speed and σ the surface tension of the resin. Due to the outer diameter of the microcatheter ($d = 2(r + h)$), the outer diameter of the microcatheter is,

$$\mathrm{d} = 2\mathrm{r}\left(1 + \frac{1.34\left(\frac{\mu V}{\sigma}\right)^{\frac{2}{3}}}{1 - 1.34\left(\frac{\mu V}{\sigma}\right)^{\frac{2}{3}}}\right)$$

Therefore, when the properties of the resin mixture remain unchanged, the outer diameter of the microcatheter is positively correlated with the pulling speed of the tungsten wire.

**TubeBot testing in BOECs model**

The bovine oviducts were obtained from a slaughterhouse (Vorwerk Podemus e.K., Dresden, Germany). Bovine oviduct epithelial cells (BOECs) were isolated and cultured according to the procedure described by Palma-Vera et al..[2] Surrounding tissues and blood vessels were removed from the oviducts, and they were cleaned with Dulbecco PBS, followed by filling the lumen with collagenase 1A (1 mg/mL in Ham's F12 medium) for 1 hr at 37°C. The mucosa was then extruded and passed through a 40 μm pore-size filter. The remaining cell clusters on the filter were washed with PBS and incubated with 0.5% trypsin/EDTA at 37°C for 10 minutes. The collected cells were either seeded on an insert or cryopreserved. A total of $2 \times 10^5$ cells were seeded onto a 0.4 μm pore-size insert (Sarstedt 24-well plate) pre-coated with rat-tail collagen I for cell culture, under conditions of 37°C, 5% $CO_2$, and 20% $O_2$. Within one week, the cells were maintained under liquid-liquid conditions using proliferation medium to promote cell division. Subsequently, the medium was removed from inside the insert to create an air-liquid interface to allow for cell differentiation. The cells were cultured for at least four weeks before experiments, with the medium being replaced twice a week.

Once a stable BOECs layer was established, it was immersed in medium and placed on the bottom of a culture dish. Cured PDMS (10:1 mass ratio) was cut into strips, placed on the BOECs layer, and covered with a coverslip, forming an in vitro model. The TubeBot was introduced into the channel of this in vitro model, and its magnetic actuation was achieved using magnets placed beneath the model, with a rotation frequency of 7 Hz and a magnetic field strength of approximately 100 mT. After several back-and-forth crawling movements within the model channel, the BOECs layer was retrieved for live/dead staining. Live cells were stained green with SYBR 14, while dead cells were stained red with Propidium Iodide (PI). The staining solution was prepared according to the kit instructions (L7011, Invitrogen). A total of 5 μL SYBR 14 was added to the channel and incubated for 10 minutes, followed by 5 μL PI for another 10 minutes. The in vitro model was then washed with pre-warmed fresh medium to remove the fluorescence staining. Images were taken using a fluorescence microscope (AxioObserver, Zeiss). SYBR 14 was visualized using a green filter (emission wavelength: 518 nm, excitation wavelength: 489 nm), while PI was visualized using a red filter (emission wavelength: 636 nm, excitation wavelength: 493 nm).

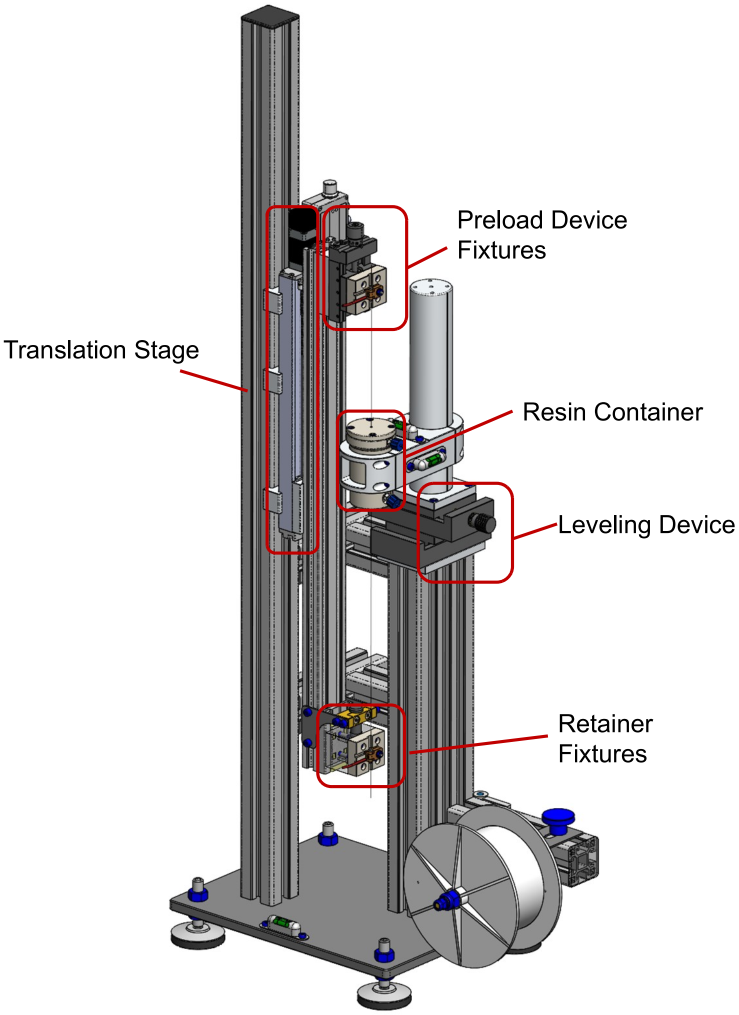


**Figure S1.** A schematic illustration of the microcatheter robot platform fabrication setup. The setup comprises key components including the resin container, preload device fixtures, leveling device, translation stage, and retainer fixtures.

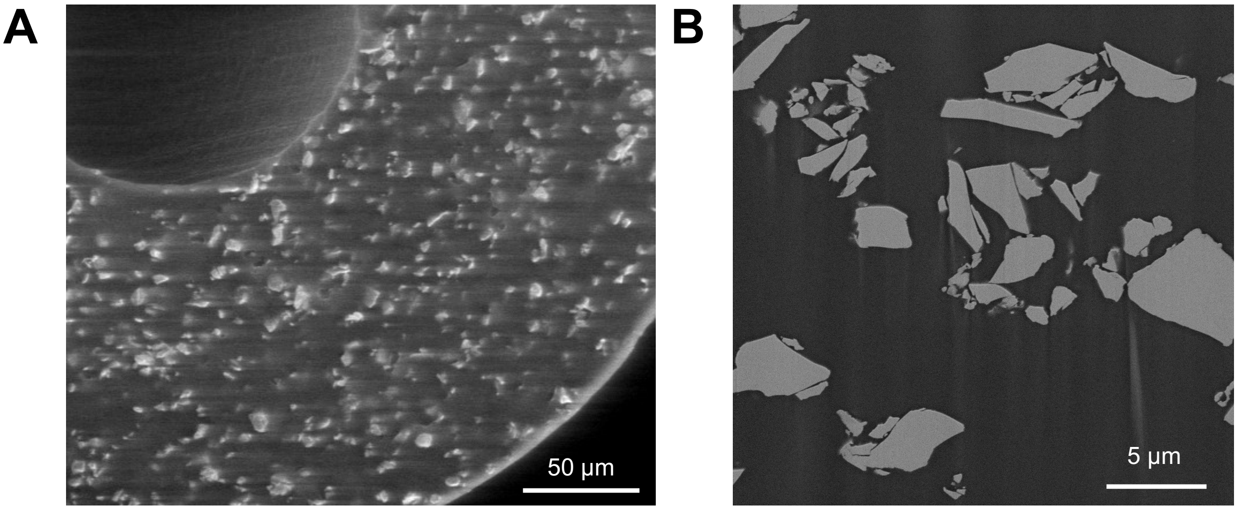


**Figure S2.** Cross-sectional scanning electron microscope (SEM) images of the magnetic microcatheter. (A) Distribution of NdFeB particles embedded in PDMS. (B) Magnified view showing individual NdFeB particle morphology.

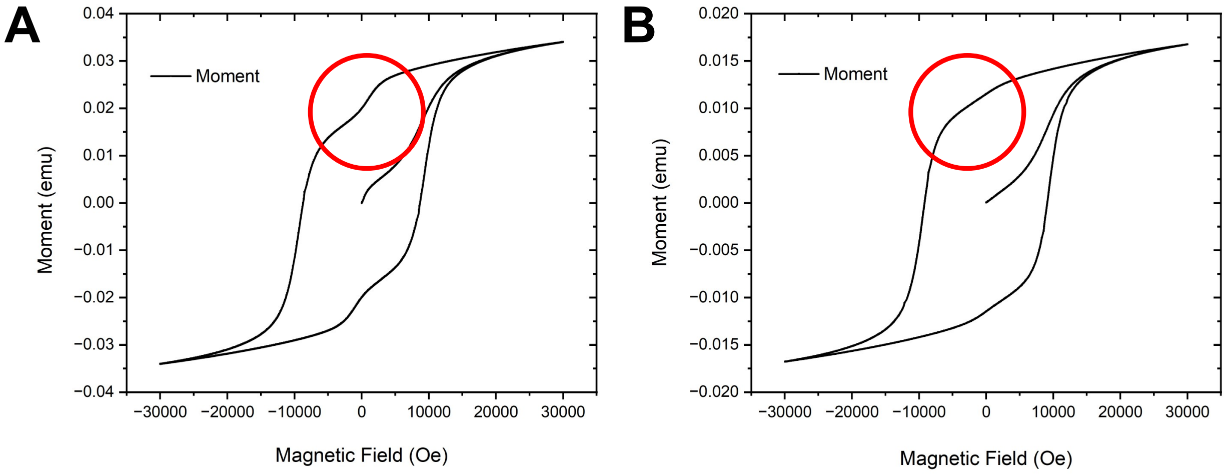


**Figure S3.** Magnetic hysteresis loops of the fabricated microcatheter samples. (A) Magnetic properties of the microcatheter cured at elevated temperatures using the fabrication setup. (B) Magnetic properties of the microcatheter cured at room temperature. The circled regions highlight differences in coercivity and remanence between the two curing methods.

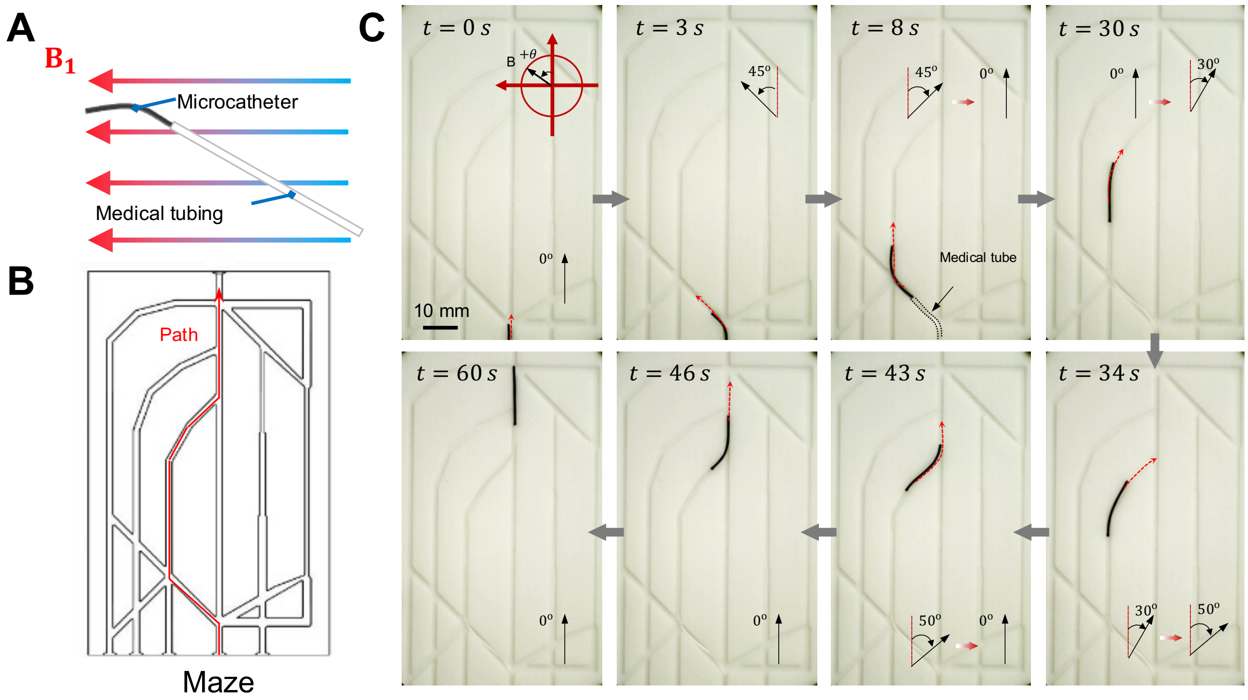


**Figure S4.** Magnetic navigation of the microcatheter robot in a 2D maze model. (A) Guiding microcatheter consisting of a microcatheter and medical tubing steered under a magnetic field. (B) The path used for navigation within the maze. (C) Sequential images illustrate the microcatheter's navigation through the maze at different times, demonstrating precise steering under magnetic control.

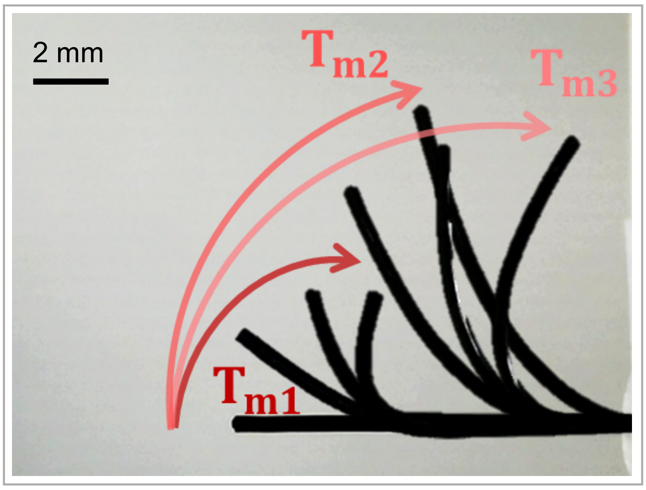


**Figure S5.** Deflection range of the guiding microcatheter with varying stiffness. The overall stiffness of the guiding microcatheter is changed by changing the length of the copper wire inserted into the working channel, and a magnetic field (0-100mT) is used to make the tip of the guiding microcatheter reach different target points in space along different trajectories.

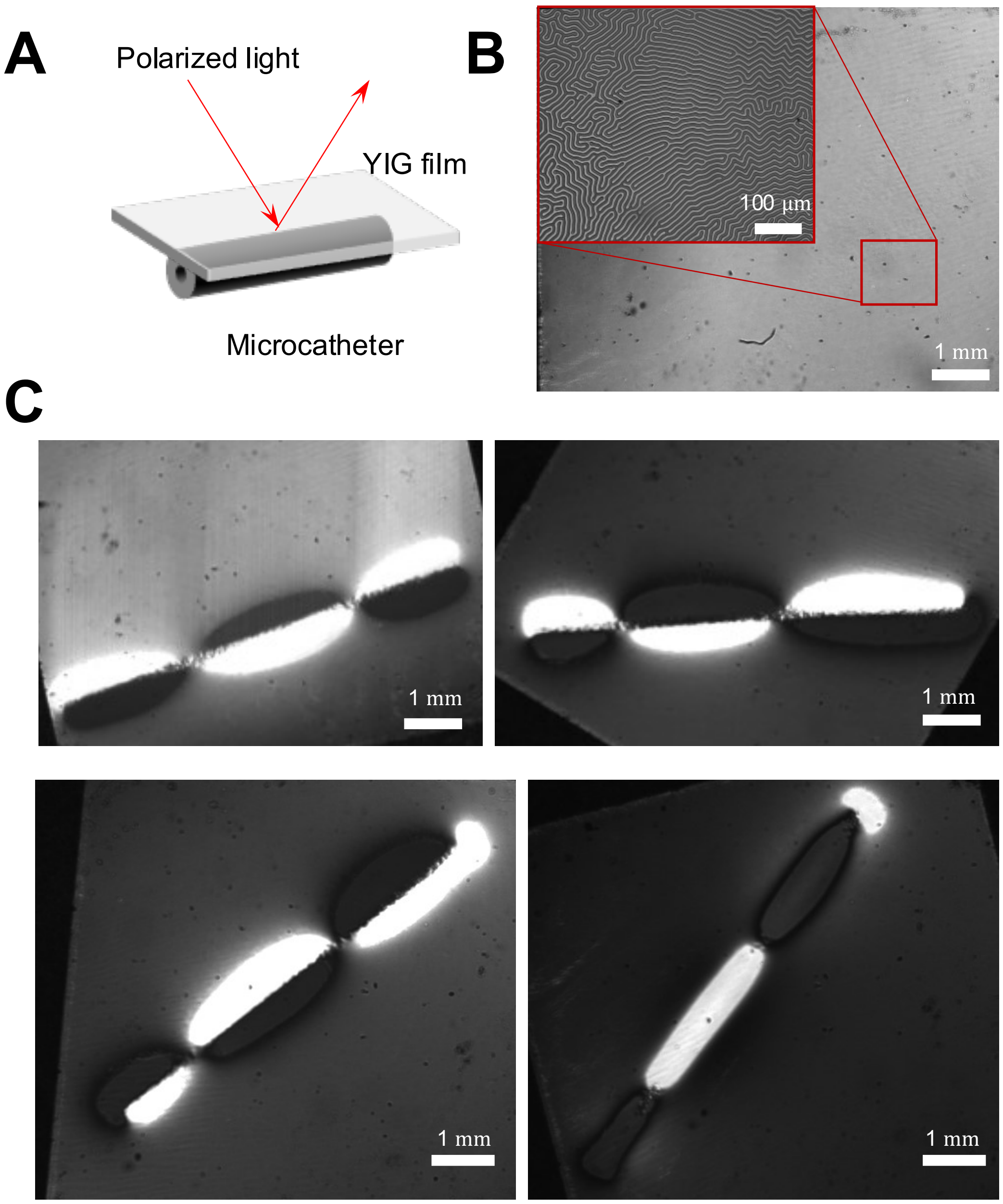


**Figure S6.** TubeBot magnetization analysis using magneto-optic Kerr effect (MOKE) microscopy. (A) A schematic experiment, utilizing magneto-optical indicator film with perpendicular anisotropy magneto-optical indicator film with perpendicular anisotropy (PMOIF) positioned on top of a cylindrical microcatheter with altering magnetization vector orientation. (B) By using a magneto-optical wide-field Kerr microscope configured for polar sensitivity under low-resolution settings—where the individual garnet domains are not visible— a pole distribution within the microcatheter can be detected. (C) PMOIF images of the magnetization distribution within the microcatheter demonstrating the presence of nodes, indicating the altering magnetization along the TubeBot.

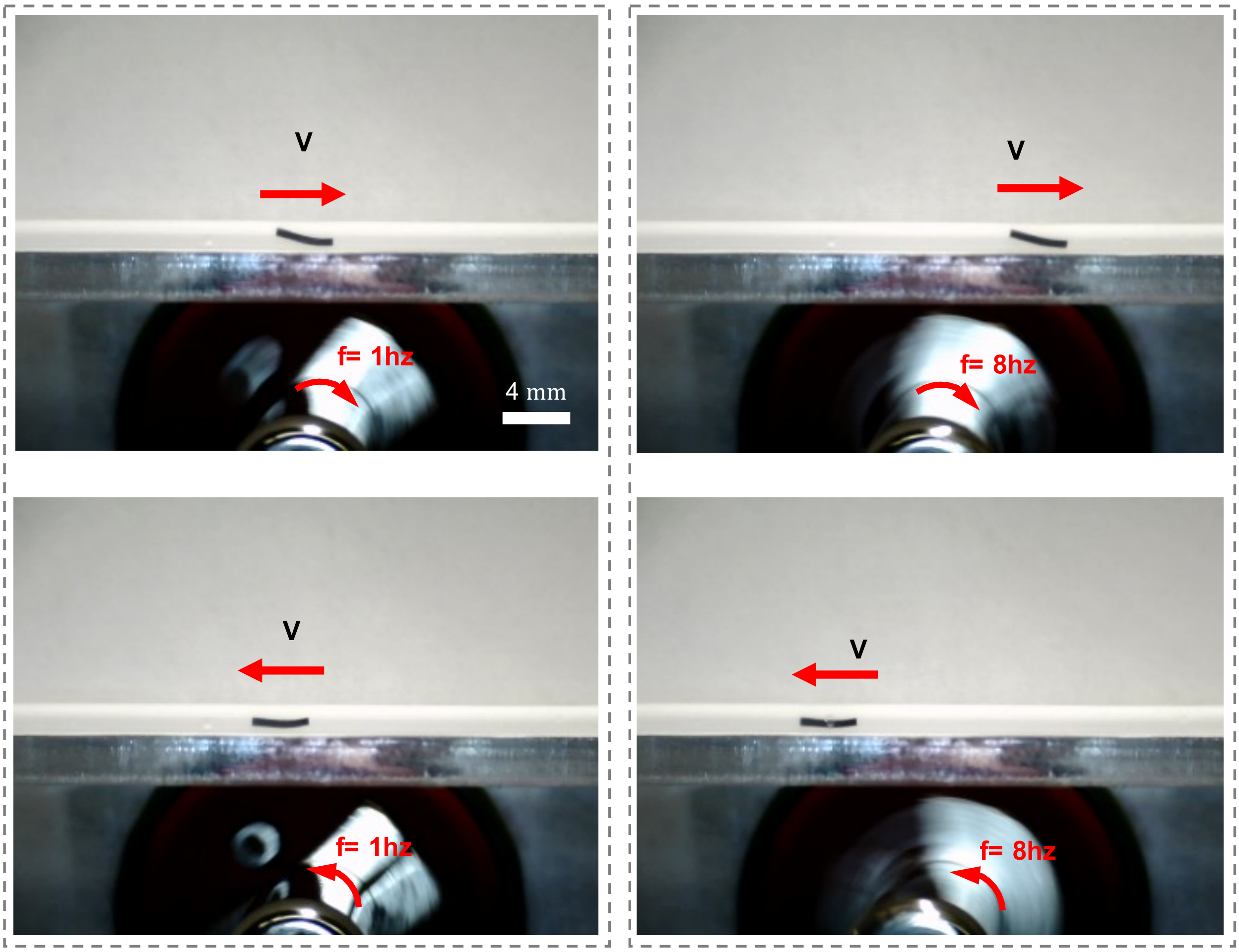


**Figure S7. TubeBot motion test.** The motion direction of TubeBot is the same as the rotation direction of the permanent magnet at different rotation frequencies.

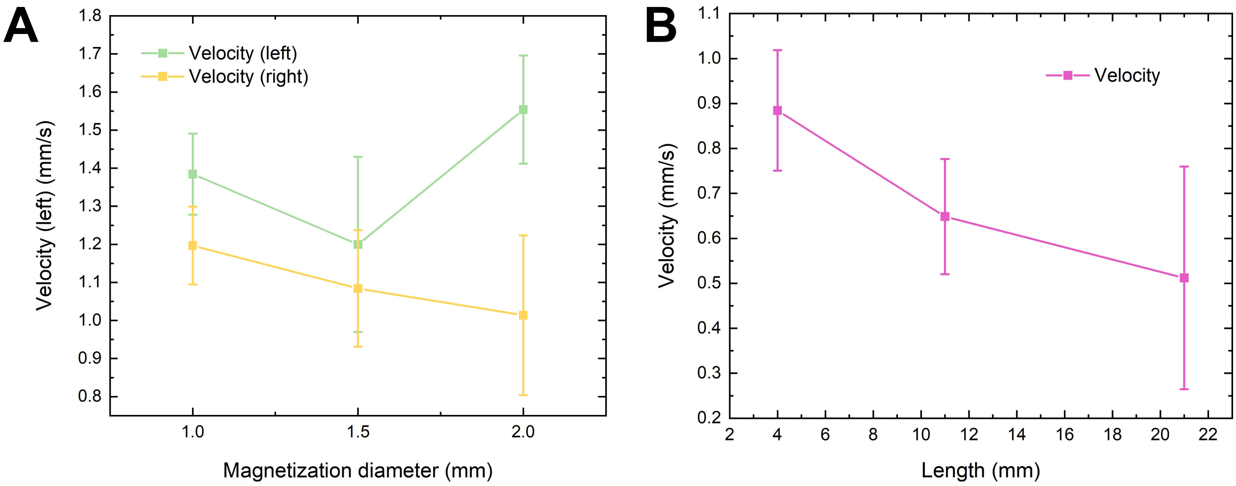


**Figure S8.** Velocity performance of the microcatheter robot. (A) Impact of different magnetization diameters, where asymmetrical magnetization results in varying speeds in different directions. (B) decreased crawling speed as microrobot length increases.

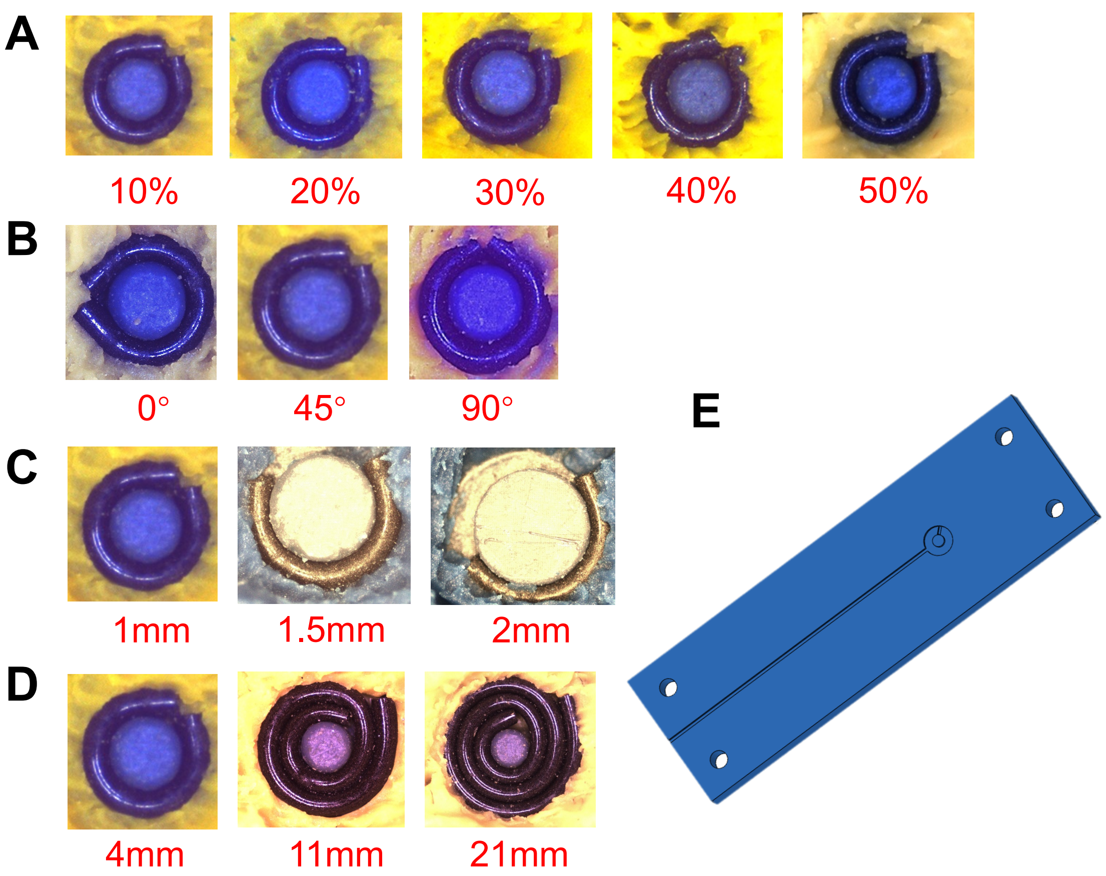


**Figure S9.** Different schemes TubeBot is magnetized. (A) Maintaining the initial magnetization angle at 45°, microcatheters with different NdFeB particle contents were magnetized. The lengths were all 4 mm. (B) Microcatheters with NdFeB particle mass fraction of 50% were magnetized at different initial magnetization angles. The lengths were all 4 mm. (C) Maintaining the initial magnetization angle at 45°, microcatheters with different magnetization diameters were magnetized with a length of 4 mm. (D) Maintaining the same magnetization diameter and initial magnetization angle, microcatheters of different lengths were magnetized. (E) The mold for magnetization.

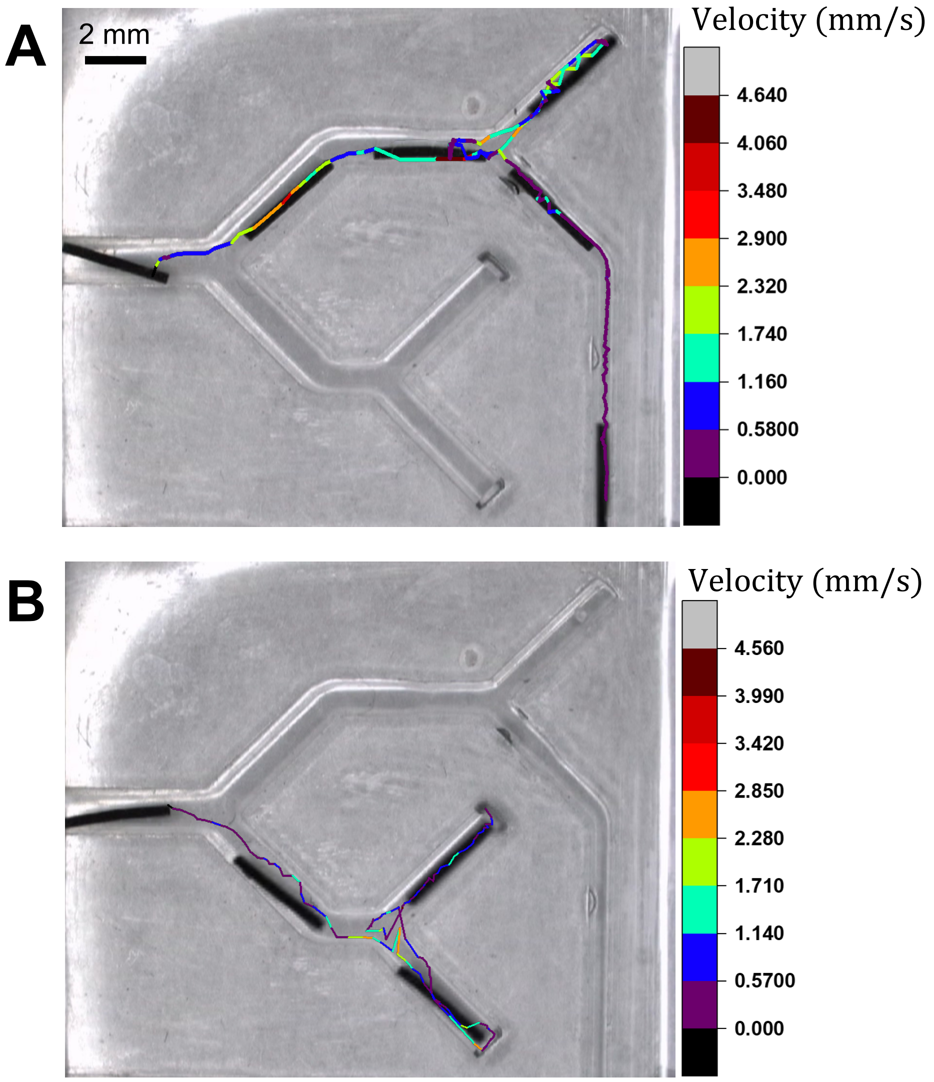


**Figure S10.** TubeBot navigating and crawling in a channel with variable diameter. Image overlays show the microrobot at different positions as it moves along the channel, with velocity tracked. (A) TubeBot navigating in path 1. (B) TubeBot navigating in path 2.

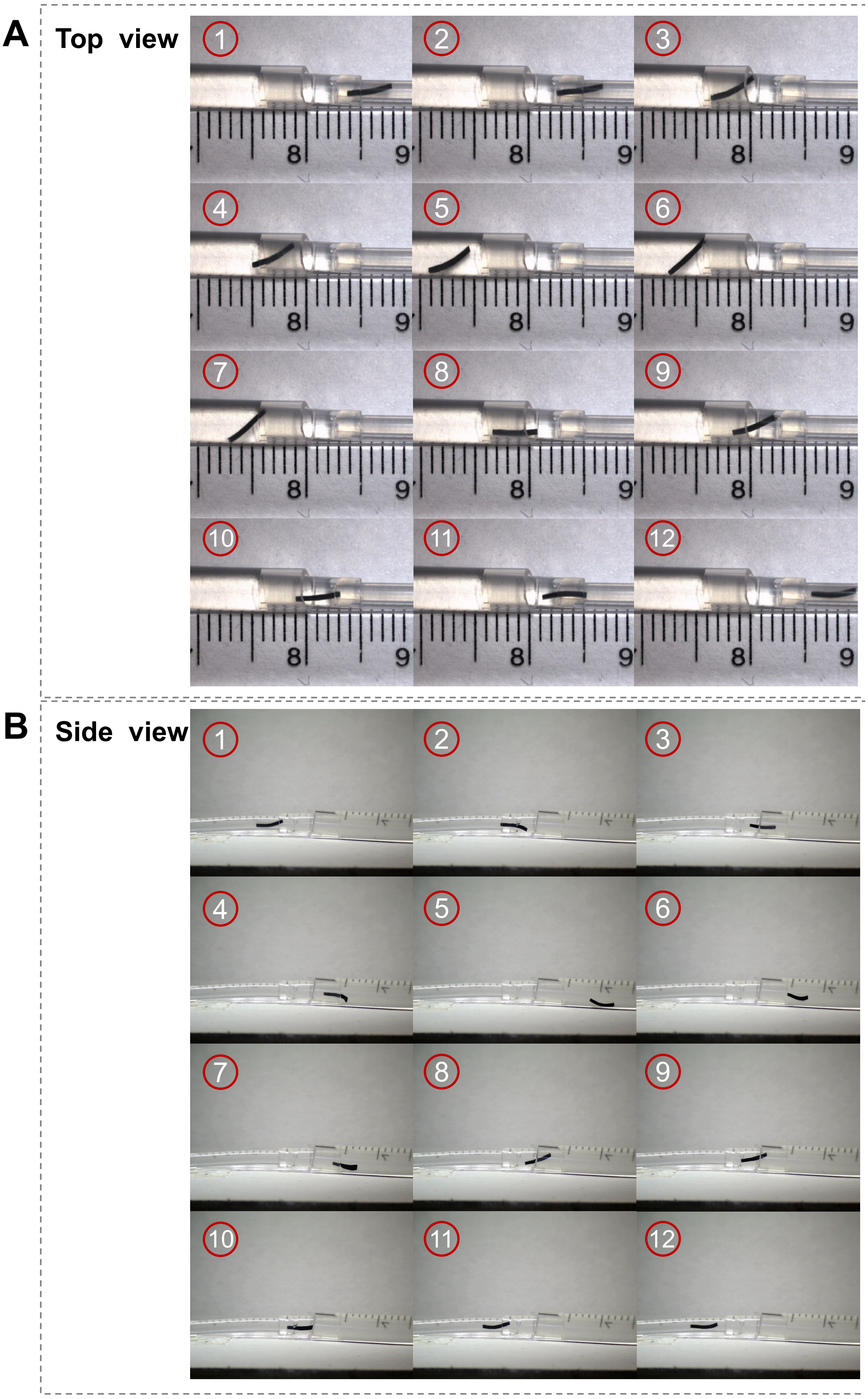


**Figure S11.** TubeBot climbs over steps in a variable diameter channel model. It transitions from a smaller diameter to a larger diameter, and then from a larger diameter to a smaller diameter. (A) Top view. (B) Side view.

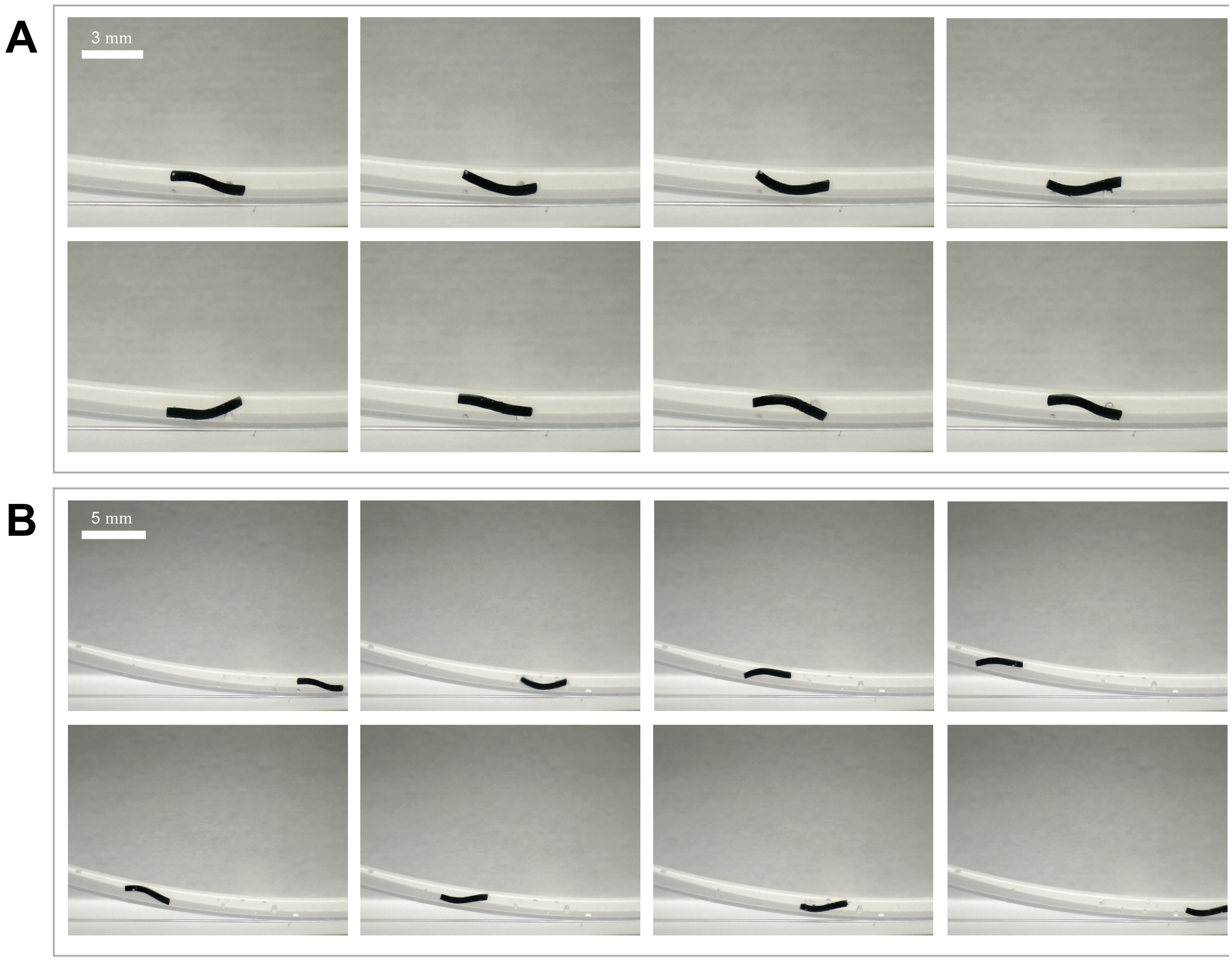


**Figure S12.** TubeBot movement test after loading cargo. (A) Deformation of alginate-filled TubeBot in a magnetic field (100mT). (B) Alginate-filled TubeBot crawling up and down in a sloping tube (8hz 100mt ID=1mm).

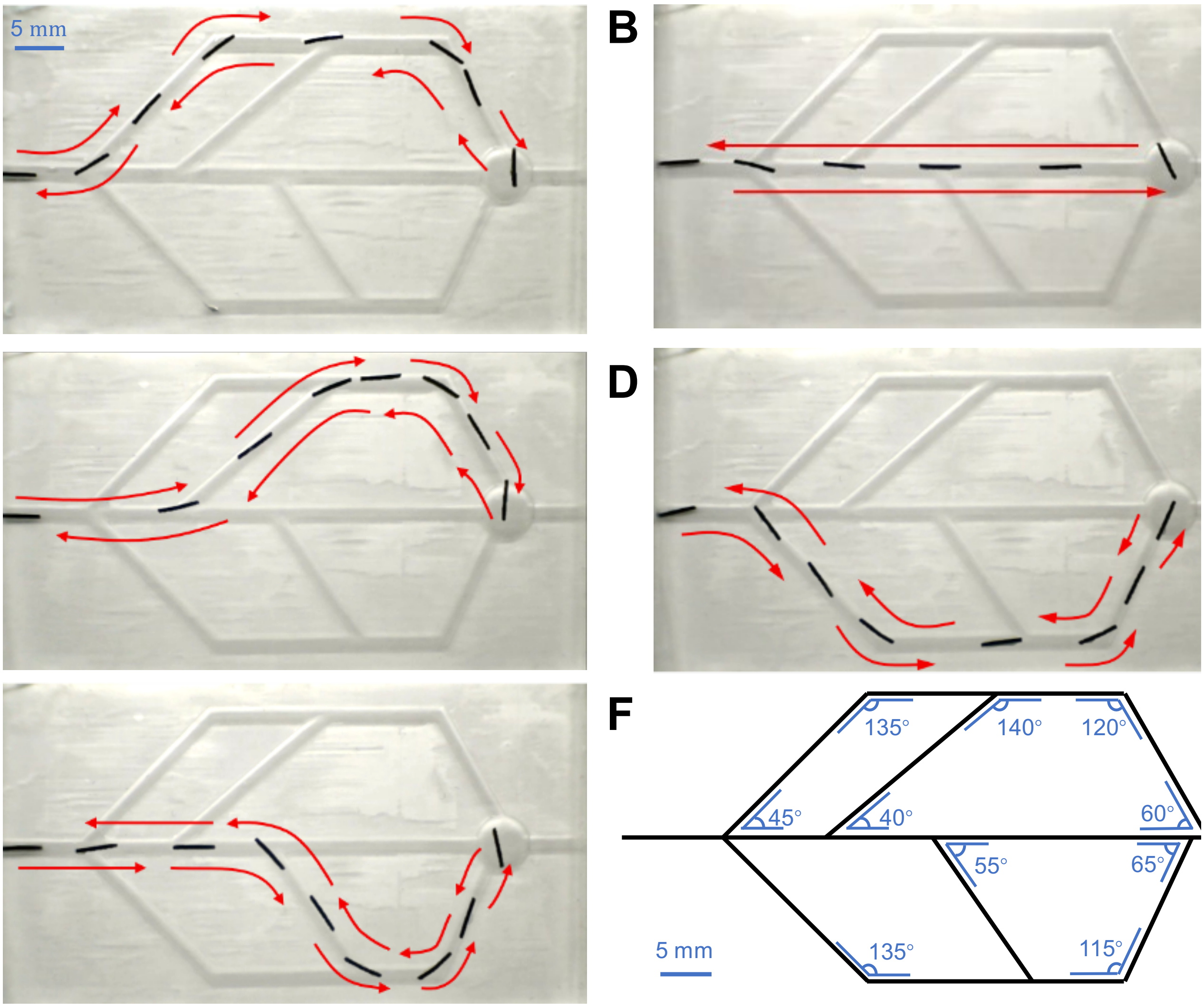


**Figure S13.** The navigation crawling of the alginate-filled TubeBot in a complex channel model. (A) The channel angle changes in sequence: 45°, 135°, 120°, 60°. (B) The channel is a straight line. (C) The channel angle changes in sequence: 40°, 140°, 120°, 60°. (D) The channel angle changes in sequence: 45°, 135°, 115°, 65°. (E) The channel angle changes in sequence: 55°, 125°, 115°, 65°. (F) The size of the complex channel model.

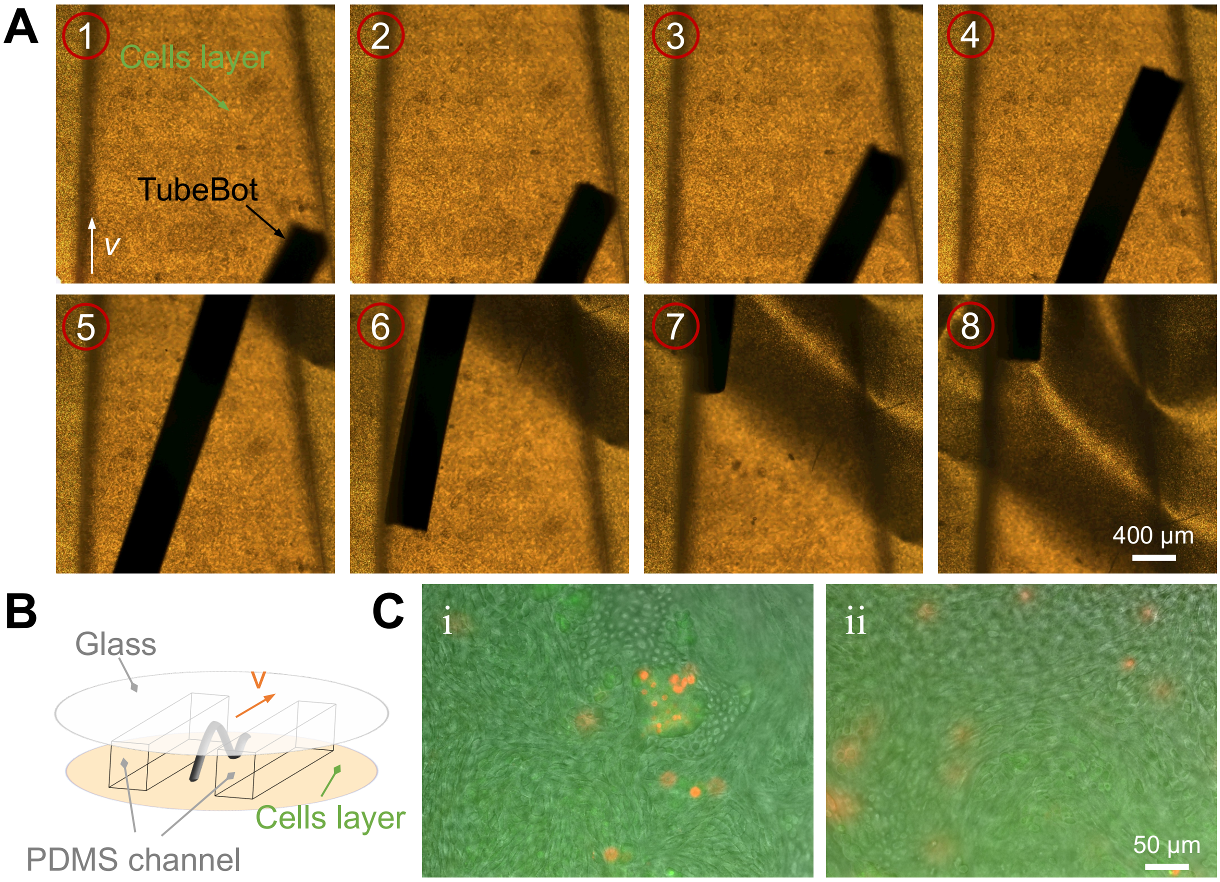


**Figure S14.** TubeBot crawls in the bovine oviduct epithelial cells layer channel model to verify the effect of movement on cell activity. (A) TubeBot crawls forward on the cell layer under an optical microscope. (B) Schematic diagram of the in vitro cell layer model. (C) Live and dead fluorescent images of the cell layer after the TubeBot crawling movement test. (i) Experimental image, (ii) control image.

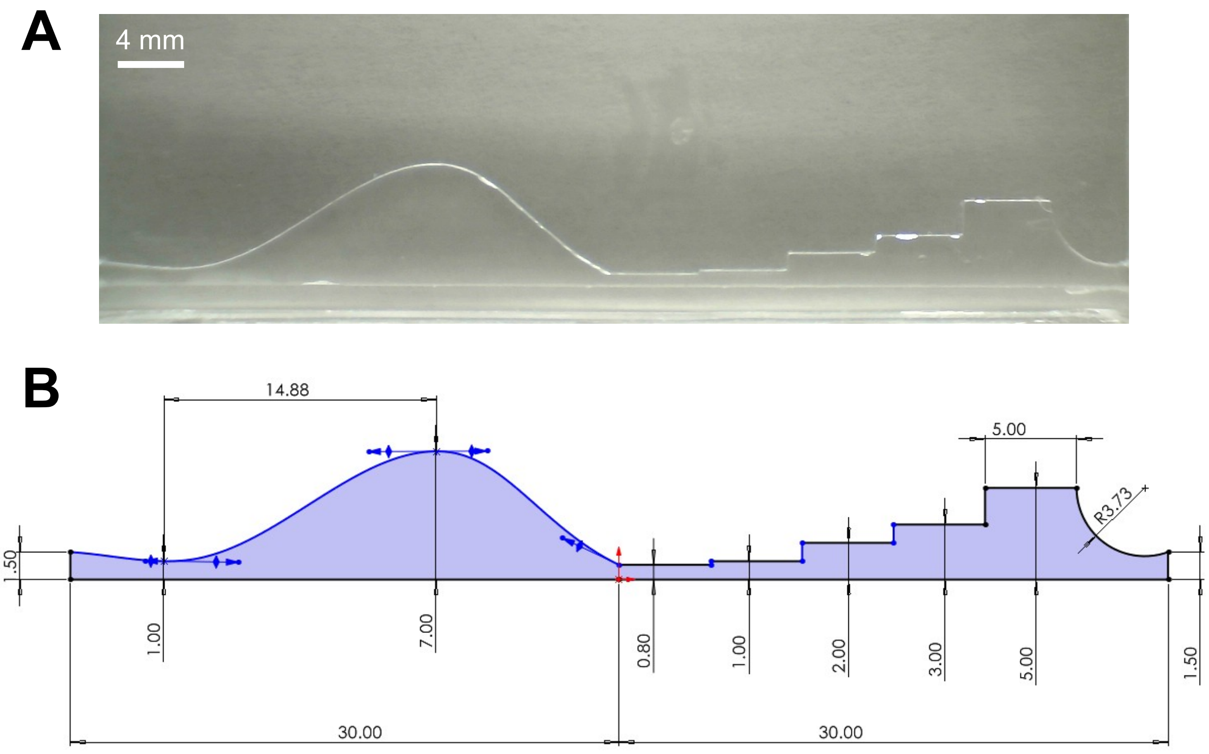


**Figure S15.** Complex channel model. (A) Image of the channel. (B) Dimensions of the channel.

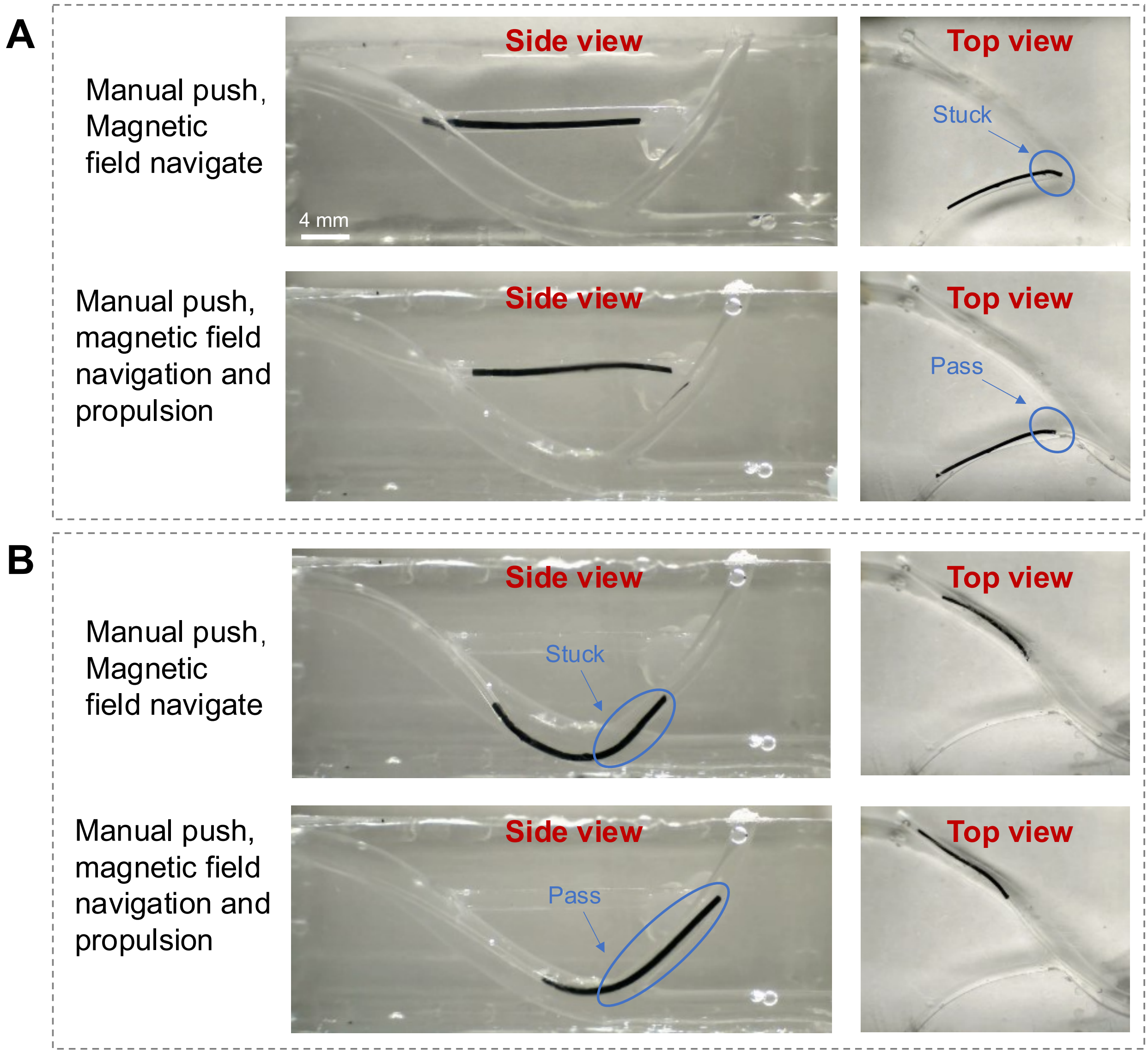


**Figure S16.** Two navigation and propulsion methods of microcatheter robot in a 3D flow channel. Comparison of manual advancement and a combination of magnetic propulsion with manual pushing. Manual push with magnetic field navigation always makes the microcatheter robot stuck in places with sharp corners or narrow passages. This does not happen with magnetic propulsion with manual push. (A) Narrow and sharp channel. (B) Upward and sharp channel.

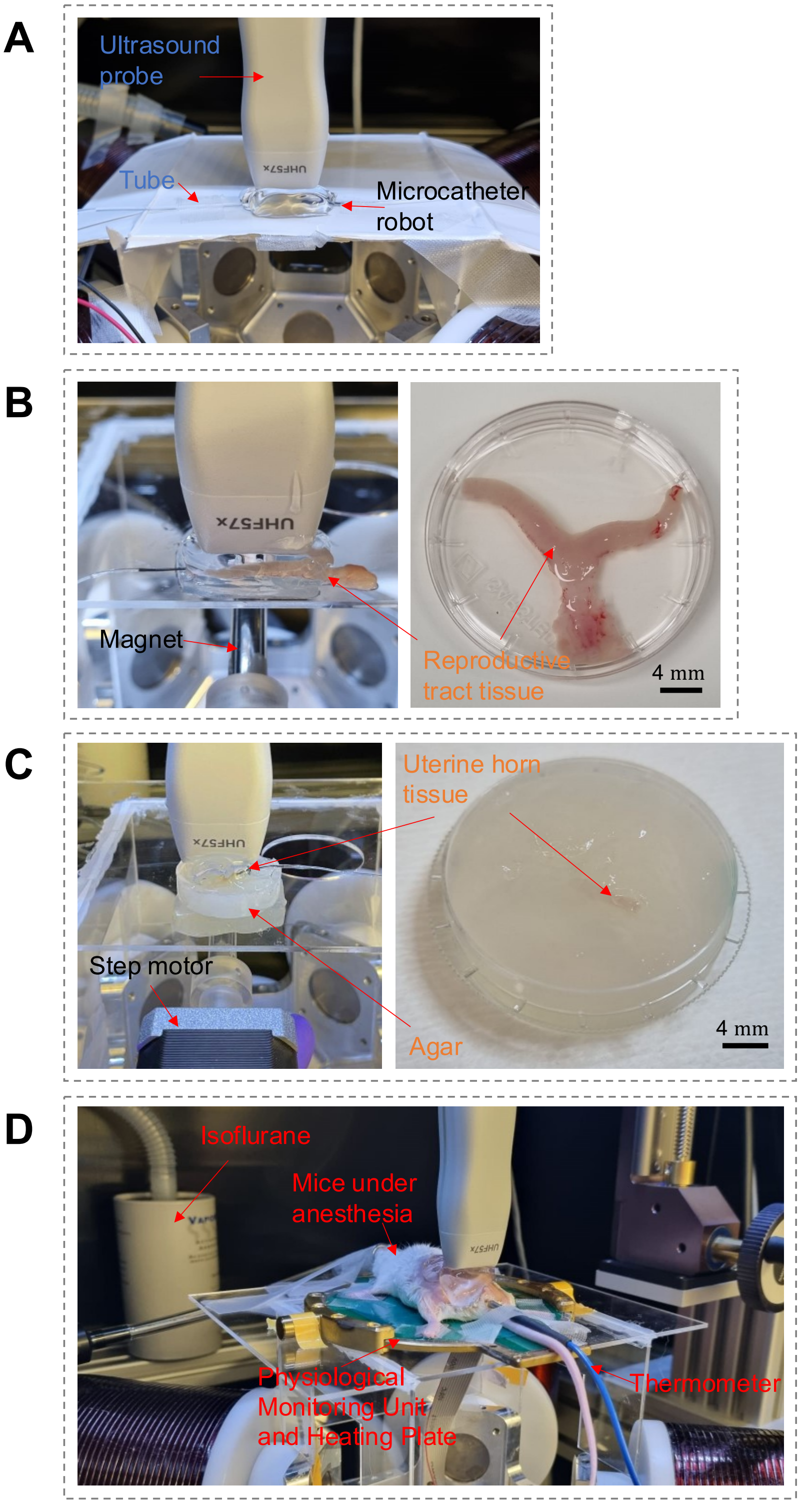


**Figure S17.** Practical setup of ex vivo and in vivo phantoms with integrated ultrasound imaging. (A) Ex vivo tubing model setup. (B) Ex vivo reproductive tract tissue model setup and reproductive tract tissue in PBS. (C) Ex vivo uterine horn tissue model setup and uterine horn tissue in agar. (D) In vivo model setup. The mice were anesthetized under a continuous supply of isoflurane, and the physiological monitoring unit monitored the physiological data of the mice and provided heating, and the body temperature was measured by a thermometer. Ultrasound probe was placed above the reproductive system.

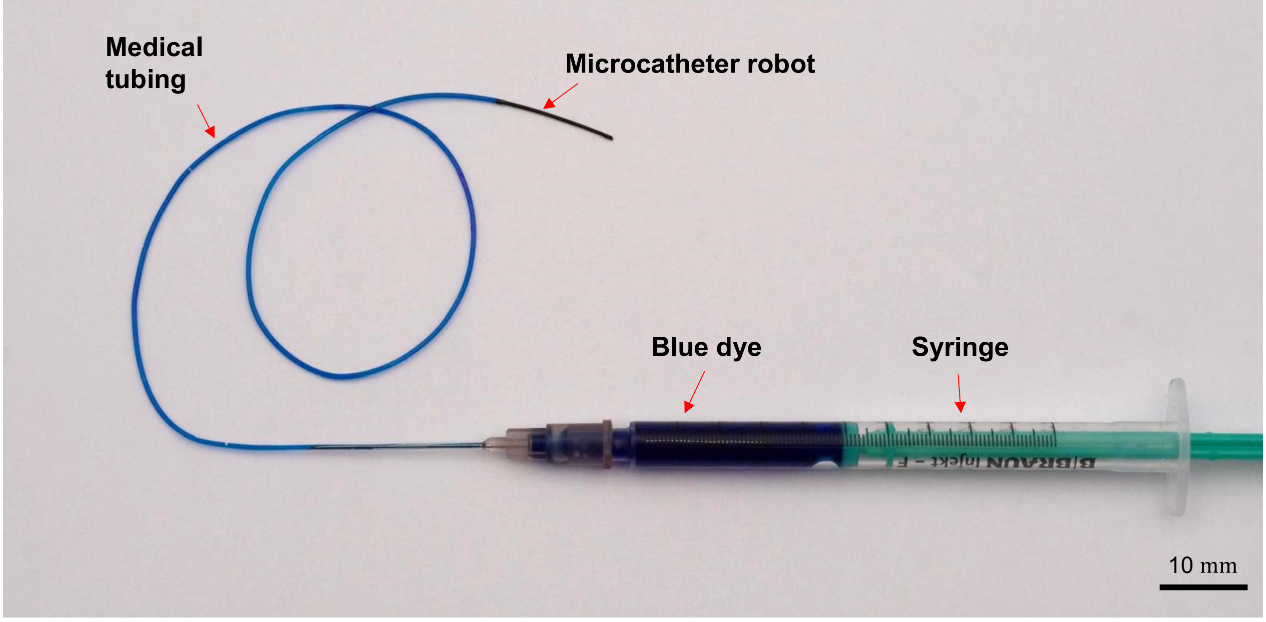


**Figure S18.** The microcatheter robot for in vivo experiments. The microcatheter robot is connected to medical tubing. One end of the medical tubing is connected to a syringe with blue dye inside.

**Table S1.** Comparison of Catheter Designs: Fabrication, Size, Navigation, and Applications.

| Catheters | Fabrication | Size [mm] | Navigation | Advancement | Adaptability | In vitro/In vivo |
|---|---|---|---|---|---|---|
| Ref. [3] | Mechanical assembly | 2 | Tendon | Mechanical push | No | In vitro |
| Ref. [4] | Mold casting | 0.9 | Hydraulic | Manual push | No | In vitro and in vivo |
| Ref. [5] | Mold casting | 1.3 | Shape memory alloy | Manual push | yes | In vitro |
| Ref. [6] | Mold casting | 1.67 | Magnet | Mechanical push | No | In vitro |
| Ref. [7] | Injection molding | 4–7 | Magnet | Mechanical push | No | Ex vivo and in vivo |
| Ref. [8] | Printing/inje ction molding | 0.5-0.8 | Magnet | Manual push | No | In vitro |
| Ref. [9] | Continuous curing | 0.35 | Magnet and flow | Flow and manual push | yes | In vitro and ex vivo |
| Ref. [10] | Injection molding | 2 | Magnet | Manual push | yes | Ex vivo and in vivo |
| Ref. [11] | Mechanical assembly | 0.7-0.8 | Magnet | Mechanical push and tip rotation | yes | Ex vivo and in vivo |
| This work | Continuous curing | 0.3-1 | Magnet | Manual push and end propulsion | yes | Ex vivo and in vivo |

**Table S2.** Comparative Analysis of Catheter Fabrication, Cost, and End Propulsion.

| Catheters | Size [mm] | Fabrication | Preparation Cost | End Propulsion |
|---|---|---|---|---|
| Ref. [12] | 2.6-6 | Mechanical assembly | High | Pneumatic Peristaltic Wave Propulsion |
| Ref. [13] | 108 | Mechanical assembly | High | Pneumatic Peristaltic Wave Propulsion |
| Ref. [14] | 25 | Mold casting and Mechanical assembly | High | Pneumatic Peristaltic Wave Propulsion |
| This work | 0.3-1 | Continuous curing | Low | Terminal undulatory crawling propulsion |
